# Poxvirus attack of anti-viral defense pathways unleashes an effector-triggered NF-κB response

**DOI:** 10.1101/2025.03.04.641538

**Authors:** Brenna C. Remick, Joshua Q. Mao, Andrew G. Manford, Ami D. Gutierrez-Jensen, Allon Wagner, Michael Rape, Grant McFadden, Masmudur M. Rahman, Moritz M. Gaidt, Russell E. Vance

## Abstract

Effector-triggered immunity (ETI) is a form of pathogen sensing that involves detection of pathogen-encoded virulence factors or “effectors”. To discover novel ETI pathways in mammals, we developed a screening approach in which individual virulence factors are expressed in human monocytes and transcriptional responses are assessed by RNA-seq. Using this approach, we identify a poxvirus effector, myxoma virus M3.1, which elicits an anti-viral NF-κB response. We find that NF-κB is unleashed by an ETI pathway that senses M3.1 attack of two anti-viral complexes: ZAP and TBK1. NF-κΒ activation occurs because the proteins inhibited by M3.1— N4BP1, ZC3H12A, and TBK1—are negative regulators of NF-κB. Our results illustrate how negative regulators can function as pathogen sensors and establish a systematic approach for the discovery of ETI pathways.

## Introduction

The mammalian innate immune system senses pathogens by detecting conserved molecular motifs called pathogen-associated molecular patterns (PAMPs) (*1*). PAMPs such as lipopolysaccharide and flagellin serve as ligands for germline-encoded receptors called pattern-recognition receptors (PRRs). Binding of a PAMP to its cognate PRR initiates immune responses referred to as PAMP-triggered immunity (PTI). Sensing of PAMPs is an effective means to differentiate self from non-self, but cannot easily discriminate between pathogens and harmless microbes, as PAMPs are present on both.

A distinguishing feature of pathogens is the expression of proteins called virulence factors or “effectors” (*2*). During infection, pathogens employ an arsenal of virulence factors to target key host factors and thereby counteract host defenses and facilitate replication. Hosts have evolved to “guard” the key pathways commonly targeted by virulence factors, such that protective immune responses are initiated when these pathways are disrupted during infection (*3, 4*). Immune sensing of virulence factor activities in host cells is referred to as effector-triggered immunity (ETI) (*3, 5*). In contrast to PTI, sensing of virulence factors (i.e., ETI) is inherently pathogen-specific, as innocuous microbes do not deliver virulence factors into host cells. The concept of ETI originated in studies of plant immunity (*3, 5*) but is increasingly recognized as a mechanism of pathogen sensing in animals as well (*6–8*). While a few examples of mammalian ETI have been characterized, it is unclear whether these cases are anecdotal examples or representative of a more ubiquitous—and perhaps underappreciated—form of pathogen detection.

Investigations of ETI are complicated by the evolutionary arms race between hosts and pathogens, which results in layers of host immune responses and pathogen counterattacks (*8, 9*). This interplay is captured by the “zig-zag” model of plant immunity (*3*), in which sensing of PAMPs by PRRs elicits a host immune response which is then blocked by pathogen virulence factors. In turn, hosts evolve mechanisms to detect virulence factors or their activities, resulting in ETI. Pathogens may then evolve additional virulence factors that suppress ETI. This ongoing cycle of adaptation results in a reciprocal pattern of susceptibility and resistance, giving rise to the “zig-zag” dynamic. Consequently, pathogens that are well-adapted to a host may fail to elicit detectable ETI responses, yet ETI pathways may still exist and confer protection in other contexts.

To circumvent this challenge to the discovery of ETI pathways, we developed a screening approach that systematically tests pathogen virulence factors for their capacity to trigger host immune responses. In this strategy, we inducibly express individual virulence factors in human BLaER1 monocytes (*10*) and then assess induction of host transcriptional responses by RNA-seq (*11*). By screening individual virulence factors, rather than whole pathogens, we enable the discovery of ETI responses that are suppressed by other virulence factors during infection. Moreover, our simplified system allows us to immediately associate a given response to a particular virulence factor.

Using the above approach, we screened a library of myxoma virus (MYXV) open reading frames (ORFs) (*12*). MYXV is a rabbit-adapted poxvirus that is notoriously lethal to European rabbits (*13*). Like other poxviruses, MYXV harbors a large, double-stranded DNA (dsDNA) genome encoding dozens of virulence factors that manipulate host cellular processes and block immune signaling pathways (*14, 15*). MYXV does not infect humans or mice, but it can infect certain human cancer cell lines (*16*). We reasoned that virulence factors from a non-adapted pathogen, such as MYXV, may be more likely to trigger ETI responses in human cells than those from a well-adapted pathogen that is under selective pressure to avoid eliciting immune responses.

## Systematic virulence factor screening identifies MYXV M3.1 as an inducer of NF-κB signaling

We employed our arrayed virulence factor screening approach to discover novel examples of ETI in human BLaER1 monocytes (Fig. 1A). Our library of virulence factors included the MYXV ORF library (*12*), as well as mCherry as a negative control and adenovirus E4ORF3, a virulence factor known to induce a very strong type I interferon (IFN) response (*17*), as a positive control. We successfully transduced BLaER1 cells and performed bulk RNA barcoding and sequencing (*11*) for 117 MYXV ORFs (Table S1).

**Fig. 1.**
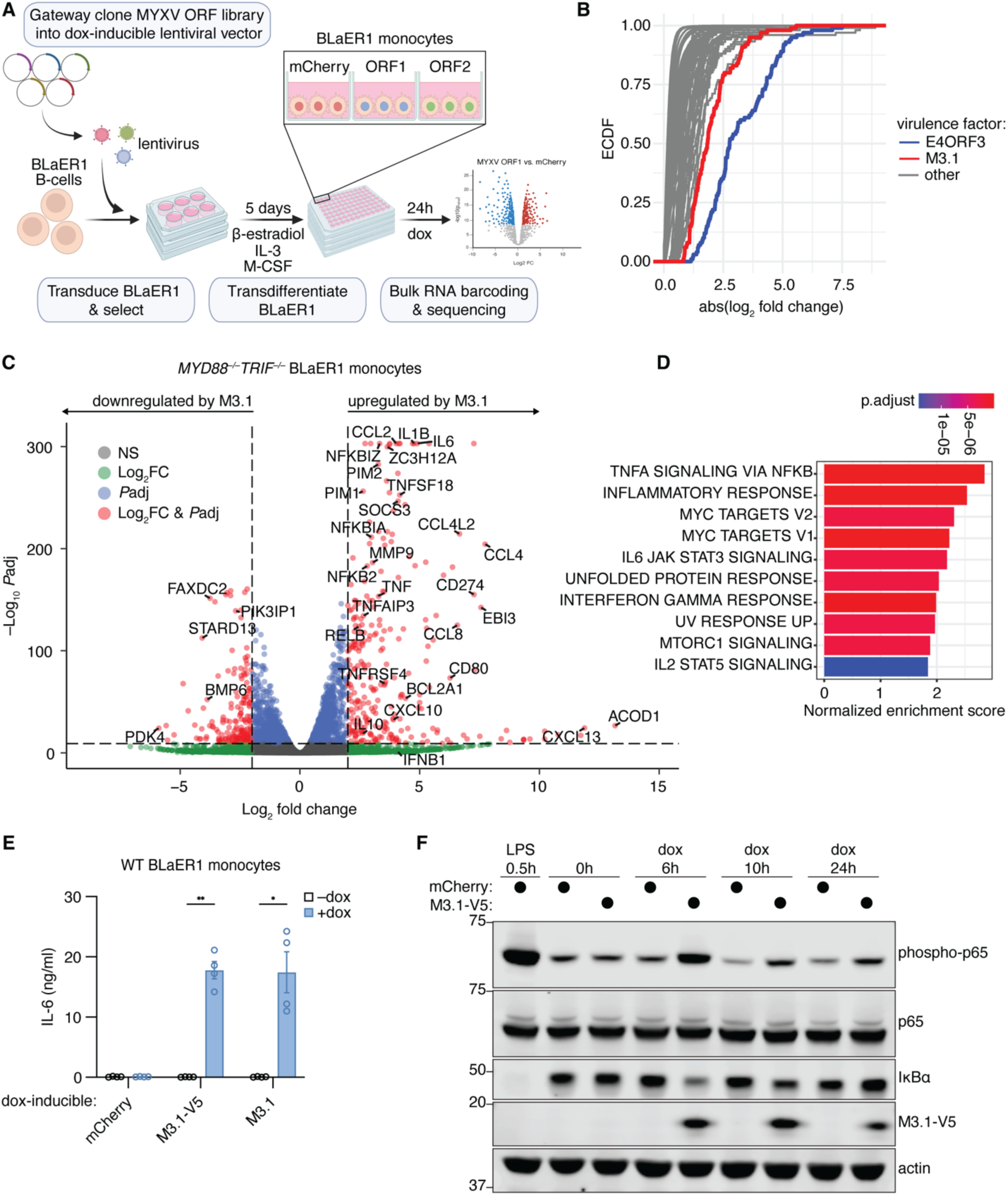
MYXV M3.1 induces NF-κB signaling in human monocytes. (**A**) Schematic of arrayed virulence factor screening approach. Created with BioRender.com. **(B)** For each ectopically expressed viral ORF, DESeq2 was used to identify differentially expressed genes compared to mCherry-expressing control samples. The empirical cumulative distribution function (ECDF) of the absolute fold change was plotted for the top 100 genes with the lowest *P*adj value for each ORF. **(C)** Volcano plot of differentially expressed genes in *MYD88^−/−^TRIF^−/−^* BLaER1 monocytes treated with doxycycline for 24 hours to induce expression of M3.1 or mCherry. Data from three independent experiments. Dashed lines indicate a log2FC cutoff of 2 and a *P*adj cutoff of 10e-10. **(D)** MSigDB hallmark gene sets enriched in M3.1-expressing *MYD88^−/−^TRIF^−/−^* BLaER1 monocytes in (C). **(E)** IL-6 secreted by BLaER1 monocytes treated with doxycycline for 24 hours. Data are mean ± SEM of four independent experiments. **(F)** BLaER1 monocytes expressing doxycycline-inducible mCherry or M3.1 were treated with doxycycline or LPS (200 ng/ml) for the indicated times. Lysates were immunoblotted as indicated. Images are representative of three independent experiments. * *P* < 0.05; ** *P* < 0.01, tested by unpaired t-test with Welch’s correction.

To identify MYXV virulence factors that elicit host transcriptional responses, we performed differential gene expression analysis on BLaER1 monocytes expressing each different viral ORF and compared these responses to samples expressing mCherry (Fig. 1B). As expected (*17*), adenovirus E4ORF3 induced a robust type I IFN response (Fig. S1), indicating that our screening strategy was performing as intended. The majority of MYXV ORFs did not elicit notable transcriptional responses. However, several MYXV ORFs induced differential gene expression, which we characterized by gene set enrichment analysis (Fig. S2). Of note, we observed MYXV M003.1 gene expression induced a strong NF-κB-like transcriptional response in BLaER1 monocytes (Fig. S2B, S3). We refer to the virulence factor encoded by the M003.1 gene as the M3.1 protein.

We confirmed the transcriptional response induced by M3.1 was independent of trace TLR ligands (e.g., LPS, lipoproteins) potentially present in the media by expressing M3.1 in *MYD88^−/−^TRIF^−/−^* BLaER1 monocytes and performing RNA-seq. Again, we observed robust induction of inflammatory cytokines and chemokines by M3.1, characteristic of an NF-κB-mediated immune response (Fig. 1C-D). We confirmed IL-6 protein induction by M3.1, which was unaffected by a C-terminal epitope tag on M3.1 (Fig. 1E). Moreover, we observed IκBα degradation and sustained p65 phosphorylation in M3.1-expressing BLaER1 monocytes (Fig. 1F), providing biochemical evidence that the NF-κB signaling pathway is activated by M3.1.

## Identification of M3.1 cellular targets by immunoprecipitation mass spectrometry

As NF-κB signaling is an established host defense response against poxviruses (*18*), we suspected that M3.1 did not evolve to activate NF-κB but instead evolved to block a different anti-viral pathway. In this scenario, activation of NF-κΒ would result from M3.1 “inadvertently” triggering a host ETI pathway (Fig. 2A). M3.1 is part of a family of poxvirus virulence factors that share a Bcl-2-like fold (*19*). While M3.1 itself has not been functionally characterized, other poxvirus Bcl-2-like proteins have been found to antagonize host anti-viral proteins and innate immune signaling pathways, typically by binding host proteins and blocking their activities (*20*).

**Fig. 2.**
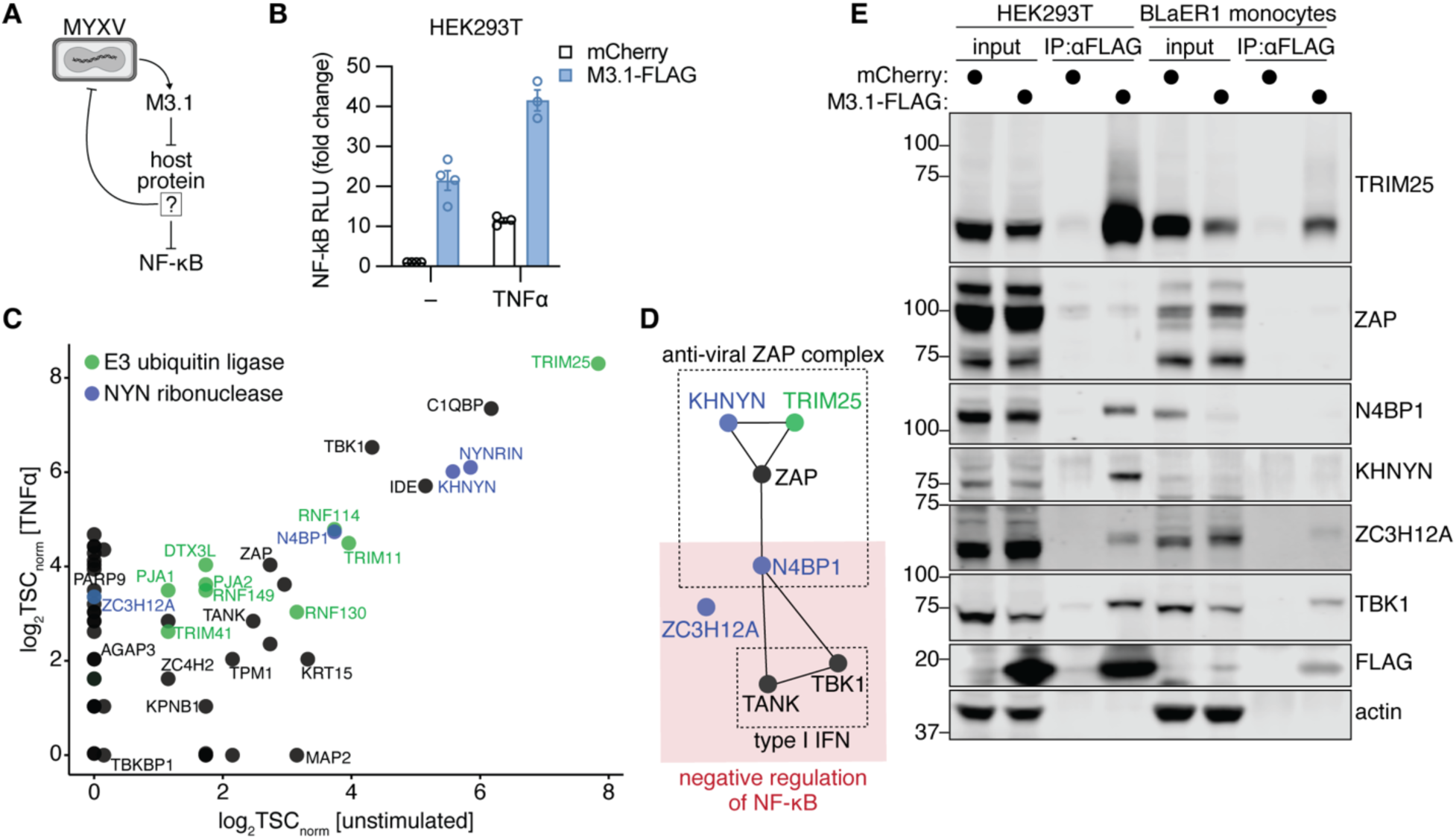
M3.1 interacts with host anti-viral factors and NF-κB inhibitors. (**A**) Hypothesized mechanism in which M3.1 inhibits an anti-viral host factor that also functions as a negative regulator of NF-κB. **(B)** A luciferase reporter was used to measure NF-κB responses in HEK293T cells transfected with M3.1-FLAG and/or stimulated with TNFα (1 ng/ml). Fold change is relative to unstimulated cells transfected with mCherry. Data are mean ± SEM of 3-4 independent experiments. **(C)** M3.1-FLAG was immunoprecipitated from HEK293T cells that were either unstimulated or treated with TNFα (10 ng/ml) for 8 hours. M3.1 binding partners were identified by mass spectrometry followed by CompPASS analysis. **(D)** String interaction network representing biochemical and/or genetic interactions between M3.1-binding proteins. **(E)** M3.1-FLAG was immunoprecipitated from HEK293T cells and BLaER1 monocytes. Co-purifying proteins were detected by immunoblotting. Images are representative of two independent experiments.

We hypothesized that M3.1 may trigger an NF-κB response by inhibiting a host anti-viral protein that also functions as a negative regulator of NF-κB (Fig. 2A). To investigate this possibility, we sought to identify M3.1-interacting proteins in human cells. We thus performed immunoprecipitation mass spectrometry (IP-MS) in HEK293T cells using FLAG-tagged M3.1 as a bait protein. To do this, we first confirmed that M3.1-FLAG is functional in HEK293T cells and is able to activate an NF-κB luciferase reporter as well as enhance NF-κB responses to TNFα stimulation (Fig. 2B). We then expressed M3.1-FLAG in unstimulated and TNFα-stimulated HEK293T cells and used IP-MS to identify M3.1-interacting proteins in both of these conditions. This approach identified many M3.1-interacting proteins, most of which were seen in both the unstimulated and TNFα-stimulated conditions (Fig. 2C).

Interestingly, several M3.1-interacting proteins have known anti-viral activities. For example, Zinc finger anti-viral protein (ZAP) restricts a broad range of RNA and DNA viruses, in some cases by binding CpG-rich viral RNA and targeting it for degradation (*21–23*). Several ZAP cofactors have been identified, including the E3 ubiquitin ligase TRIM25 (*24, 25*) and the NYN ribonucleases KHNYN and N4BP1 (*26–28*), all of which were also identified by IP-MS as proteins that interact with M3.1. Moreover, additional NYN ribonucleases, including the KHNYN and N4BP1 paralog NYNRIN, as well as ZC3H12A (MCPIP1, REGNASE-1) also interacted with M3.1. Notably, N4BP1 and ZC3H12A are known negative regulators of NF-κB signaling (*29–31*). The IP-MS results also indicated that M3.1 interacts with TBK1 and its adaptor TANK. TBK1 is primarily known for its role in inducing type I IFN, a critical host anti-viral transcriptional response. Recently, however, suppression of NF-κB by N4BP1 has been shown to involve TBK1 and TANK (*32*). Thus, many of the proteins we observed to interact with M3.1 are functionally linked to each other and to key anti-viral signaling pathways (Fig. 2D).

We used coimmunoprecipitation to validate M3.1 interactions with TRIM25, NYN ribonucleases, and TBK1 in both HEK293T cells and BLaER1 monocytes (Fig. 2E). While we observed interactions between M3.1 and ZAP cofactors TRIM25, KHNYN, and N4BP1, we were unable to detect a robust interaction between M3.1 and ZAP itself, suggesting a weak or indirect interaction that was unstable in our coimmunoprecipitation conditions. We also observed reduced protein levels for M3.1-interacting proteins, particularly N4BP1, in BLaER1 lysates and, to a lesser extent, in HEK293T lysates, raising the possibility that M3.1 may promote degradation of its binding partners. Together, these results indicate that M3.1 interacts both with anti-viral proteins and negative regulators of NF-κB in human cells.

## MYXV employs M3.1 to escape restriction by the ZAP complex and to suppress TBK1-mediated type I IFN signaling

We hypothesized that M3.1 evolved primarily to block the anti-viral ZAP complex and/or TBK1-mediated type I IFN signaling, but in doing so, unleashes NF-κB signaling, possibly though suppression of NF-κB inhibitors N4BP1 and ZC3H12A. While ZAP has been found to restrict vaccinia virus (VACV) (*33*), a poxvirus related to MYXV, it is unknown whether ZAP restricts MYXV. To test if ZAP restricts MYXV replication in human cells, and if M3.1 is required to block ZAP anti-viral activity, we generated M3.1-deficient MYXV (vMyx-ΔM003.1) and infected both HEK293T cells and BLaER1 monocytes. MYXV replicated poorly in BLaER1 monocytes (Fig. S4), so we focused on HEK293T cells. We observed that, while MYXV replicates efficiently in HEK293T cells, vMyx-ΔM003.1 was severely attenuated (Fig. 3A-B). Thus, M3.1 promotes MYXV replication.

**Fig. 3.**
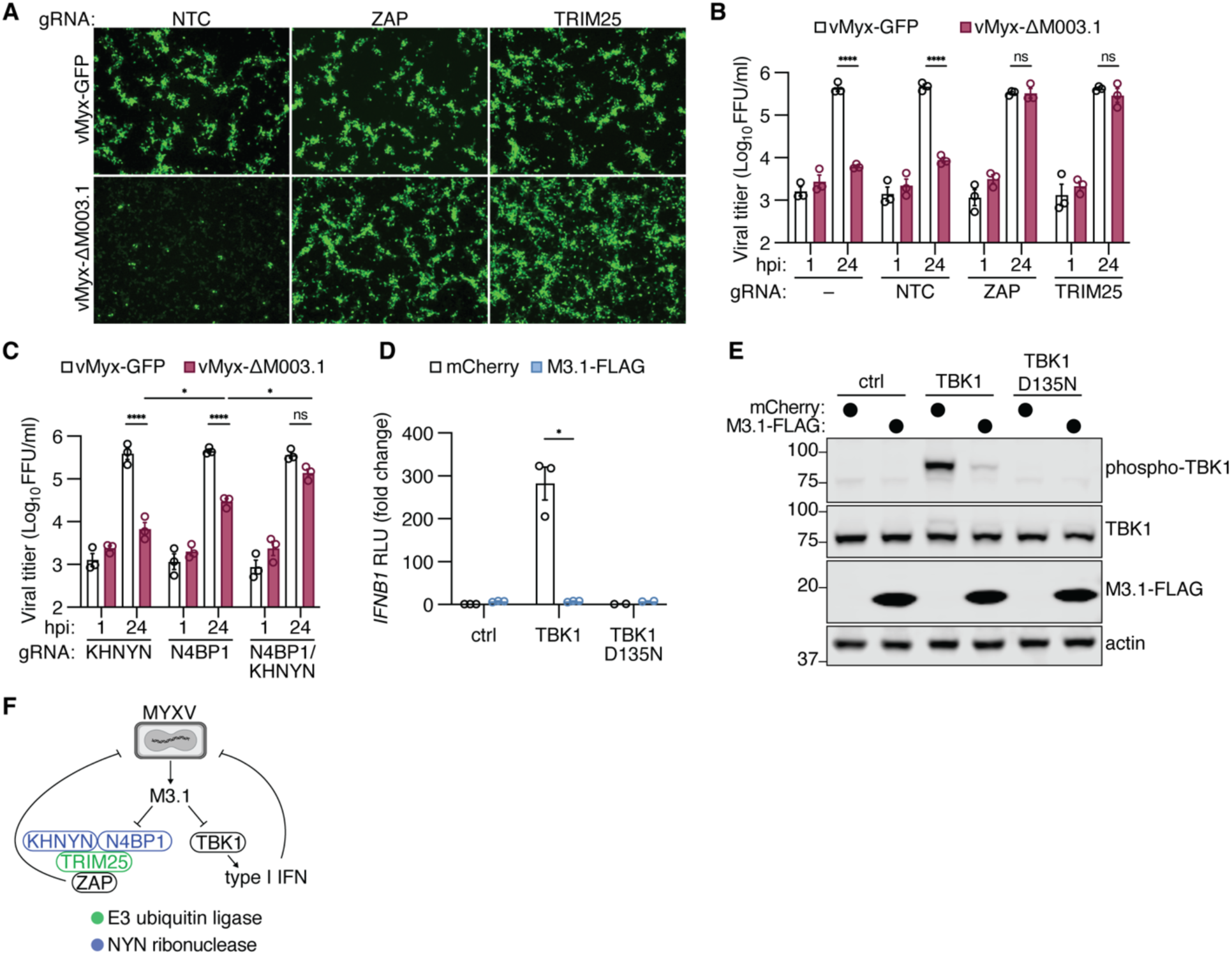
M3.1 blocks ZAP complex anti-viral activity and inhibits TBK1-mediated type I IFN induction. (**A**) HEK293T cells nucleofected with Cas9 and two guides targeting the indicated gene or a non-targeting control (NTC) were infected with GFP-expressing MYXV (MOI = 1) for 24 hours. Images were taken using an inverted microscope at 4x magnification. Images are representative of three independent experiments. **(B-C)** MYXV replication in HEK293T cells infected with an MOI of 1. Viral progeny were quantified at the indicated time points. Data are mean ± SEM of 3 independent experiments. **(D)** A luciferase reporter was used to measure *IFNB1* induction in HEK293T cells co-transfected with wild-type or kinase-dead (D135N) TBK1 (25 ng) and M3.1-FLAG (50 ng). Fold change is relative to cells transfected with mCherry. Data are mean ± SEM of 2-3 independent experiments. **(E)** Lysates from (D) were immunoblotted as indicated. Images are representative of one experiment. **(F)** Model in which M3.1 inhibits two key anti-viral pathways: the ZAP complex and TBK1-mediated type I IFN signaling. * *P* < 0.05; **** *P* < 0.0001; ns = not significant, tested by 3-way ANOVA of log-normalized data with Tukey’s post-hoc test (B-C) or by unpaired t-test with Welch’s correction (D).

Genetic deletion of ZAP, or its cofactor TRIM25, in HEK293T cells by Cas9 ribonucleoprotein (RNP) nucleofection (*34*) (validation in Fig. S5), fully rescued vMyx-ΔM003.1 replication, indicating that M3.1 is required to block ZAP anti-viral activity during MYXV infection (Fig. 3A-B). While the NYN ribonuclease KHNYN is required for ZAP-mediated restriction of retroviruses, KHNYN is dispensable for ZAP anti-viral activity in other contexts (*21*). Deletion of *KHNYN* failed to rescue vMyx-ΔM003.1 replication, indicating that KHNYN is not required for ZAP-mediated restriction of MYXV in HEK293T cells (Fig. 3C). Recent work has shown that KHNYN and its paralog N4BP1 have partially redundant functions in the ZAP complex (*26*), so we hypothesized that N4BP1 may be compensating for *KHNYN* deficiency. We observed that *N4BP1* deletion modestly increased vMyx-ΔM003.1 replication and that co-deletion of *N4BP1* and *KHNYN* largely rescued vMyx-ΔM003.1 replication (Fig. 3C). Thus, N4BP1 and KHNYN redundantly promote ZAP-mediated restriction of MYXV, and MYXV employs M3.1 to escape this restriction.

While our results suggest that a major function of M3.1 is to block ZAP anti-viral activity, we considered the possibility that M3.1 may have additional activities that support virus infection. Since we observed an interaction between M3.1 and TBK1, we also tested whether M3.1 affected type I IFN signaling. Overexpression of TBK1 in HEK293T cells activates an *IFNB1* promoter luciferase reporter in a kinase-dependent manner (*35, 36*) (Fig. 3D). Co-expression of M3.1 completely blocked TBK1-mediated *IFNB1* reporter activation (Fig. 3D) and suppressed TBK1 phosphorylation (Fig. 3E). Together, our results support a model in which M3.1 has two pro-viral functions: (1) inhibition of ZAP complex anti-viral activity, and (2) suppression of TBK1-mediated type I IFN signaling (Fig. 3F).

## M3.1 triggers an NF-kB response by blocking N4BP1, ZC3H12A, and TBK1

We next wanted to understand how M3.1 activates NF-κB signaling. Since the ZAP complex and type I IFN signaling are important anti-viral pathways, we hypothesized that hosts have evolved to guard these pathways, such that their disruption by a virulence factor, such as M3.1, triggers a compensatory NF-κB response. To explain how this occurs, we hypothesized that the anti-viral factors targeted by M3.1 may possess an additional function of repressing NF-κB. Indeed, of the M3.1-interacting proteins we identified, N4BP1, ZC3H12A, and TBK1 are all reported to inhibit NF-κB responses (*29–32, 37*). We thus predicted that genetic deletion of these M3.1 target(s) might induce NF-κB signaling, phenocopying M3.1 expression.

To test this idea, we used Cas9-RNP nucleofection to genetically disrupt M3.1-interacting partners in BLaER1 cells that also express doxycycline-inducible M3.1 (*34*) (validation in Fig. S6). We then measured IL-6 production as a readout of NF-κB activation with and without M3.1 expression (Fig. 4A). In BlaER1 monocytes, deletion of *ZC3H12A* alone induced IL-6 production, but this response was further elevated by M3.1 expression, suggesting that additional M3.1 targets are functional NF-κB repressors that are inactivated by M3.1 (Fig. 4A). Notably, co-deletion of NYN ribonucleases *ZC3H12A* and *N4BP1* drove significant IL-6 production, which was not further enhanced by M3.1 expression (Fig. 4A). Deletion of the related NYN ribonuclease *KHNYN* did not affect IL-6 production. Likewise, deletion of *TBK1*, its homolog *IKKε*, or their adaptor *TANK* did not elicit IL-6 production or affect NF-κB induction by M3.1.

**Fig. 4.**
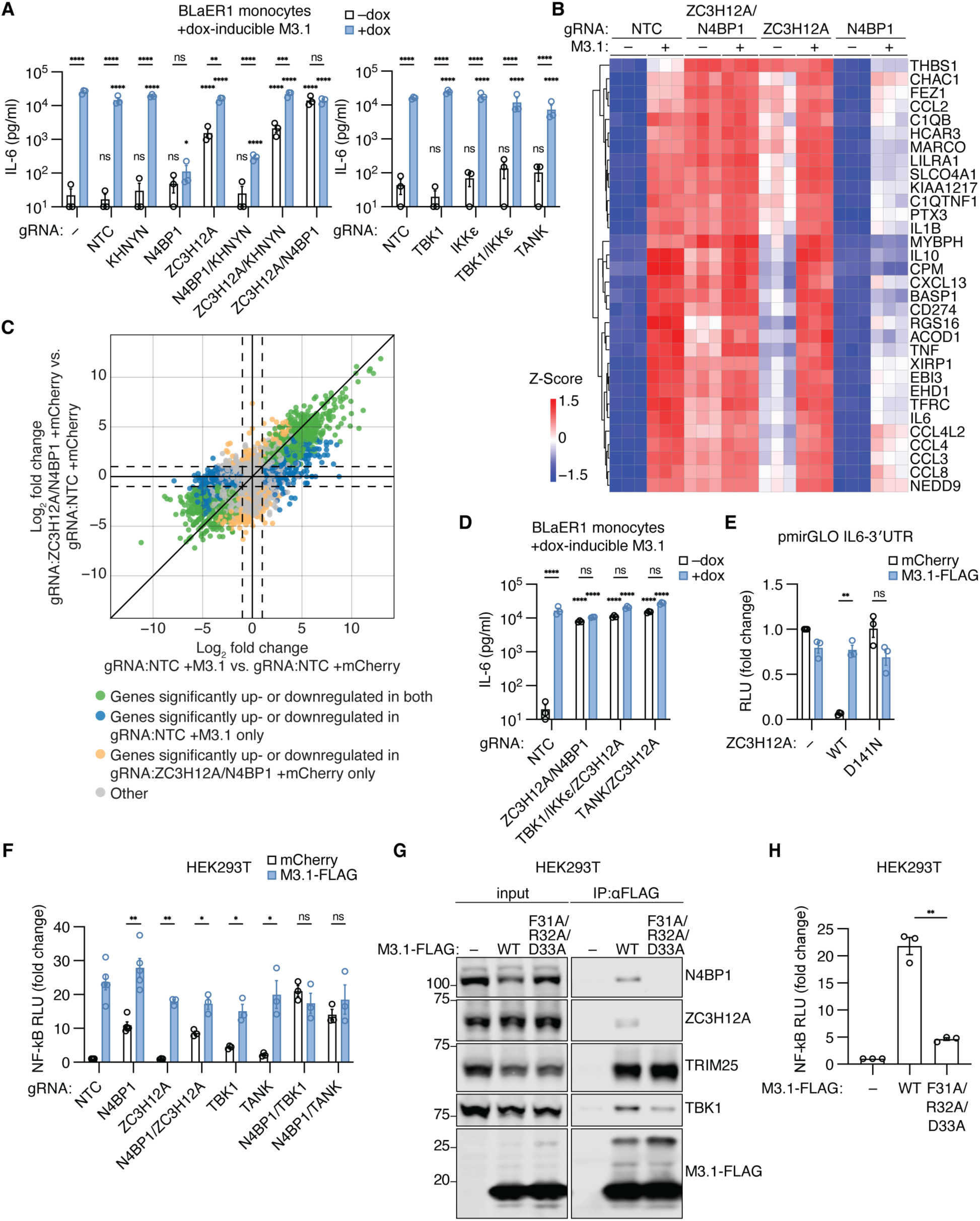
M3.1 unleashes NF-κB signaling by blocking N4BP1, ZC3H12A, and TBK1. **(A, D)** BLaER1 cells expressing doxycycline-inducible M3.1 were nucleofected with Cas9 and two gRNAs targeting the indicated gene. IL-6 secreted by BLaER1 monocytes was measured by ELISA after 24 hours of doxycycline treatment. Data are mean ± SEM of three independent experiments. **(B)** Heatmap analysis of gene expression by BLaER1 monocytes as detected by RNA-seq in three independent experiments. Selected genes are the top 30 genes by variance, as well as known NF-κB targets TNFα and IL-6. **(C)** Plot comparing differential gene expression among BLaER1 monocytes expressing M3.1 and *ZC3H12A/N4BP1*-deficient BLaER1 monocytes expressing mCherry. Dashed lines indicate a log_2_FC cutoff of –1 and 1. **(E)** A pmirGLO dual-luciferase plasmid containing the *IL-6* 3′-UTR was used to measure ZC3H12A ribonuclease activity. HEK293T cells were transfected with pmirGLO IL6-3′UTR (50 ng), ZC3H12A (25 ng), and M3.1 (50 ng). Luciferase activity was measured after 24 hours. Fold change is relative to mCherry-expressing control cells. Data are mean ± SEM of three independent experiments. **(F)** Cas9-RNP nucleofection was used to disrupt the indicated genes in HEK293T cells, and NF-κB induction by M3.1 was measured using a luciferase reporter. Fold change is relative to mCherry-expressing control cells. Data are mean ± SEM of 3-5 independent experiments. **(G)** FLAG-tagged wild-type (WT) or mutant (F31A/R32A/D33A) M3.1 was immunoprecipitated from HEK293T cells, and binding partners were assessed by immunoblotting. Images are representative of three independent experiments. **(H)** A luciferase reporter was used to measure NF-κB induction in HEK293T cells transfected with M3.1 variants (50 ng). Data are mean ± SEM of three independent experiments. * *P* < 0.05; ** *P* < 0.01; *** *P* < 0.001; **** *P* < 0.0001; ns = not significant, tested by 2-way ANOVA with Šídák’s post-hoc test on log-normalized data (A, D) or unpaired t-test with Welch’s correction (E-F, H). Comparisons were made to untreated control cells unless otherwise stated (A, D).

To more broadly characterize the transcriptional profile of *ZC3H12A/N4BP1*-deficient BLaER1 monocytes, we performed RNA-seq of BLaER1 monocytes lacking one or both factors, with and without M3.1 expression (validation in Fig. S6F). Consistent with our ELISA data, *ZC3H12A/N4BP1*-deficent cells phenocopied M3.1-expressing control cells, and expression of M3.1 had little effect on the transcriptional profile of *ZC3H12A/N4BP1*-deficent cells (Fig. 4B). Moreover, there was a striking correlation between the transcriptional responses of M3.1-expressing cells and *ZC3H12A/N4BP1*-deficent cells (Fig. 4C). Together, these data support a model where M3.1 unleashes NF-κB signaling in BLaER1 monocytes by inhibiting two negative regulators of NF-κB signaling: ZC3H12A and N4BP1.

The finding that deletion of both *ZC3H12A* and *N4BP1* is necessary to fully recapitulate the effects of M3.1 is consistent with previous reports which found that mutation of *ZC3H12A* alone or *N4BP1* alone has modest or negligible effects on NF-κB induction at baseline (*29–31, 38*). Puzzlingly, we also observed that the NF-κB response induced by M3.1 was reduced in *N4BP1*-deficient cells (Fig. 4A-B). This result highlights a complex multilayered interaction between M3.1 and N4BP1. Despite this complexity, our data strongly suggest that M3.1 induces NF-κB in BlaER1 cells through inhibition of ZC3H12A and N4BP1.

Although N4BP1 and ZC3H12A both contain a NYN ribonuclease domain, they suppress NF-κB responses through distinct mechanisms. NF-κB suppression by N4BP1 is independent of its ribonuclease activity and instead depends on the ubiquitin-binding activity of N4BP1 (*31, 32*). A recent study found that deletion of *Tbk1* and *Ikke*, or their adaptor *Tank*, in mouse bone marrow-derived macrophages (BMDMs) phenocopies *N4bp1* deficiency, supporting a model where N4BP1 acts in concert with TBK1/IKKε to limit the duration of IKKα/β signaling (*32*). Consistent with these data, deletion of *ZC3H12A* in *TBK1/IKKε*-or *TANK*-deficient BLaER1 monocytes phenocopied *ZC3H12A/N4BP1*-deficient monocytes, and similarly, these monocytes did not exhibit further IL-6 production upon M3.1 expression (Fig. 4D).

ZC3H12A suppresses NF-κB responses post-transcriptionally by degrading proinflammatory mRNA transcripts via its NYN ribonuclease domain (*30*). The targets of ZC3H12A, including the *IL-6* mRNA, are recognized via a 3′ UTR stem-loop structure (*30*). To test if M3.1 blocks ZC3H12A ribonuclease activity, we adapted a luciferase reporter assay where the luciferase mRNA harbors the *IL-6* 3′ UTR (*30*). As expected, expression of ZC3H12A reduced luciferase activity while expression of a RNase catalytic mutant (D141N) did not (Fig. 4E). Co-expression of M3.1 completely restored luciferase activity, indicating that M3.1 blocks ZC3H12A ribonuclease activity.

Our data from BLaER1 monocytes suggest that M3.1 triggers an NF-κB response by inhibiting the NYN ribonucleases N4BP1 and ZC3H12A. We next asked whether M3.1 activates NF-κB signaling in HEK293T cells through a similar mechanism. Interestingly, we observed that N4BP1-deficiency alone induced an NF-κB luciferase reporter 10-fold in HEK293T cells (Fig. 4F). Unlike in BLaER1 monocytes, deletion of *ZC3H12A* was not required to mimic the effects of M3.1, and loss of *ZC3H12A* had no effect on luciferase activity. This finding is consistent with the known mechanism of ZC3H12A, which suppresses the NF-κB response post-transcriptionally by degrading proinflammatory mRNAs, an activity that would not be expected to affect an NF-κB transcriptional reporter. Notably, M3.1 expression in *N4BP1*-deficient HEK293T cells further enhanced NF-κB responses, indicating that M3.1 has activities in addition to N4BP1 inhibition that contribute to NF-κB activation in these cells. Because TBK1 is involved in NF-κB inhibition by N4BP1 (*32*) and is inhibited by M3.1 (Fig. 3D-E), we hypothesized that the increased luciferase activity in M3.1-expressing *N4BP1*-deficient HEK293T cells may be due to TBK1 inhibition by M3.1. Notably, deletion of either *TBK1* or its adaptor *TANK* in *N4BP1*-deficient HEK293T cells resulted in increased NF-κB activation which was not further enhanced by M3.1 (Fig. 4F). Together, these data support a model where M3.1 unleashes NF-κB signaling in HEK293T cells by disrupting N4BP1/TBK1-mediated NF-κB suppression.

Our data from both BLaER1 monocytes and HEK293T cells reveal a critical role for inhibition of NYN ribonucleases in the M3.1-mediated NF-κB response. Using a series of N4BP1 domain mutant constructs (Fig. S7A), we observed that the NYN ribonuclease domain is necessary and sufficient for M3.1 to interact with N4BP1 (Fig. S7B). To test the importance of the interaction between M3.1 and NYN ribonucleases, we used AlphaFold 3 (*39*) to predict the structures of the complexes formed by M3.1 with N4BP1 and ZC3H12A (Fig. S8). In agreement with our coimmunoprecipitation data (Fig. S7B), M3.1 interacts with the NYN ribonuclease domains of N4BP1 and ZC3H12A in the predicted structures (Fig. S8). We observed that the F31/R32/D33 residues of M3.1 formed an interface with the NYN domains of both N4BP1 and ZC3H12A (Fig. S8) and that mutation of these resides to alanine ablated the interaction between M3.1 and NYN ribonucleases while largely maintaining interactions with other binding partners TRIM25 and TBK1 (Fig. 4G). Importantly, this M3.1 mutant exhibited significantly reduced NF-κB activation in both HEK293T cells and BLaER1 monocytes (Fig. 4H, S9). Together, these data indicate that the interaction between M3.1 and NYN ribonucleases N4BP1 and ZC3H12A is required for NF-κB induction. We propose a model where M3.1 unleashes an NF-κB response by inhibiting negative regulators of NF-κB signaling: N4BP1, ZC3H12A, and TBK (Fig. S10A).

## TRIM25 promotes M3.1-mediated NF-κB signaling

In addition to interacting with NYN ribonucleases and TBK1, M3.1 also strongly interacts with TRIM25, an E3 ubiquitin ligase and ZAP cofactor. While we did not observe a role for ZAP in M3.1-mediated NF-κB signaling, we found that TRIM25 deficiency unexpectedly reduced NF-κB induction by M3.1 in both BLaER1 monocytes and HEK293T cells (Fig. S11A-B). Moreover, overexpression of TRIM25 in HEK293T cells further enhanced NF-κB induction by M3.1 (Fig. S11C), suggesting that TRIM25 promotes M3.1-mediated NF-κB signaling. We hypothesized that TRIM25 might provide a necessary scaffolding function to promote the stability of M3.1 complexes, or alternatively, that TRIM25 ubiquitin ligase activity might be necessary for NF-κB induction by M3.1. To test the role of TRIM25 catalytic activity, we generated two TRIM25 mutants with disrupted ubiquitin ligase activity: C50S/C53S, which blocks zinc finger coordination, and R54P, which prevents the interaction with E2 conjugating enzymes (*25, 40*). First, we tested if the mutations affected the interaction between TRIM25 and M3.1. The interaction with M3.1 was reduced by the R54P mutation and completely blocked by the C50S/C53S mutation (Fig. S11D), suggesting that M3.1 interacts with the TRIM25 RING domain, a model that is supported by the AlphaFold 3-predicted structure of the complex (Fig. S11E). Despite its reduced interaction with M3.1, the TRIM25 R54P catalytic-dead mutant was nevertheless capable of rescuing NF-κB responses in TRIM25-deficent HEK293T cells (Fig. S11C). As expected, the TRIM25 C50S/C53S mutant, which does not interact with M3.1, was unable to rescue NF-κB induction. Together, these data support a model where the interaction between M3.1 and TRIM25 promotes M3.1-mediated NF-κB responses, independent of TRIM25 ubiquitin ligase activity. One possible model would be that TRIM25 performs a necessary scaffolding function that promotes the activity of M3.1 to disrupt the function of its other target proteins.

## M3.1 orthologs are a rapidly evolving family of virulence factors that modulate NF-κB signaling

We next tested if M3.1 orthologs from other poxviruses induced NF-κB signaling. M3.1 orthologs from closely related viruses rabbit fibroma virus, swinepox virus, and lumpy skin disease activated an NF-κB luciferase reporter to varying degrees when expressed in HEK293T cells (Fig. 5A-B). In contrast, M3.1 orthologs from more distantly related poxviruses failed to induce NF-κB. B14, the VACV (Western Reserve strain) ortholog of M3.1, has been reported to suppress NF-κB responses by directly binding and inhibiting IKKβ (*41*). This finding is interesting in light of our results showing that M3.1 interacts with and inhibits TBK1 (Fig. 2-3), an IKKβ homolog. We confirmed that VACV B14, as well as its cowpox virus ortholog, blocked NF-κB induction by TNFα in HEK293T cells (Fig. 5C). VACV A52, a Bcl-2-like virulence factor reported to activate MAPK signaling (*42*), failed to induce or suppress NF-κB signaling (Fig. 5A-C). Since M3.1 orthologs from rabbit fibroma virus and swinepox virus induced strong NF-κB responses, we tested if these orthologs activated NF-κB through a similar mechanism to M3.1. In contrast to expression of M3.1, expression of the rabbit fibroma virus and swinepox virus orthologs further elevated NF-κB induction in *N4BP1/TBK1*-deficient HEK293T cells, suggesting that these orthologs have different and/or additional activities compared to M3.1 (Fig. 5D). Overall, these results highlight the rapidly evolving functions of the Bcl-2 family of poxvirus virulence factors, consistent with the hypothesis that these proteins are engaged in an evolutionary arms race with their hosts (*9*).

**Fig. 5.**
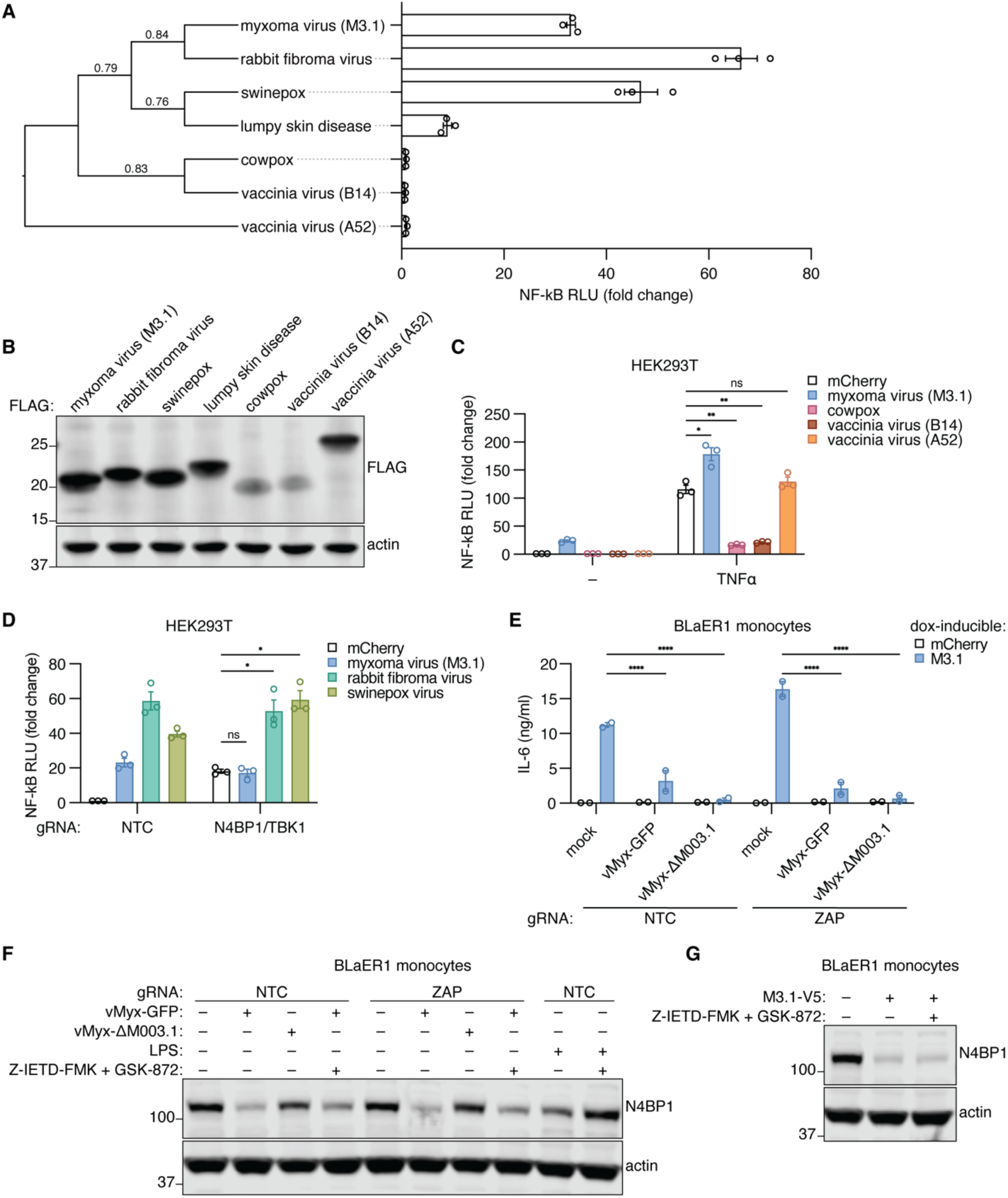
Poxvirus virulence factors modulate NF-κB signaling. **(A)** A phylogeny was built using orthologous M3.1 protein sequences from poxviruses. NF-κB induction in HEK293T cells was measured for each ortholog using a luciferase reporter. Fold change is relative to mCherry-expressing control cells. Data are mean ± SEM of three independent experiments. **(B)** Lysates in (A) were immunoblotted as indicated. Images are representative of three independent experiments. **(C)** A luciferase reporter was used to measure NF-κB responses in HEK293T cells transfected with M3.1 and its orthologs with and without TNFα (100 ng/ml). Fold change is relative to unstimulated cells transfected with mCherry. Data are mean ± SEM of 3 independent experiments. **(D)** Cas9-RNP nucleofection was used to disrupt the indicated genes in HEK293T cells, and NF-κB induction by M3.1 and its orthologs (50 ng) was measured using a luciferase reporter. Fold change is relative to mCherry-expressing control cells. Data are mean ± SEM of three independent experiments. **(E)** BLaER1 monocytes expressing doxycycline-inducible mCherry or M3.1 were treated with doxycycline and infected with MYXV (MOI = 3) for 24 hours, and secreted IL-6 was measured by ELISA. Data are mean ± SEM of 2 independent experiments. **(F)** BLaER1 monocytes were either stimulated with LPS (200 ng/ml) or infected with MYXV (MOI = 3) for 24 hours in the presence of caspase-8 (10 μM Z-IETD-FMK) and RIPK3 (3 μM GSK-872) inhibition or DMSO control. Lysates were immunoblotted as indicated. Images are representative of two independent experiments. **(G)** BLaER1 monocytes expressing doxycycline-inducible M3.1 or mCherry were treated with doxycycline for 24 hours in the presence of caspase-8 (10 μM Z-IETD-FMK) and RIPK3 (3 μM GSK-872) inhibition or DMSO control. Lysates were immunoblotted as indicated. Images are representative of two independent experiments. * *P* < 0.05; ** *P* < 0.01; **** *P* < 0.0001; ns = not significant, tested by unpaired t-test with Welch’s correction (C-D) or by 2-way ANOVA with Šídák’s post-hoc test (E).

## M3.1 targets N4BP1 during MYXV infection

Last, we sought to test whether M3.1 activates NF-κB signaling during MYXV infection. MYXV is known to encode multiple virulence factors that suppress NF-κB responses (*43–45*). Not surprisingly, we failed to detect NF-κB activation by MYXV in both HEK293T cells (Fig. S12A) and BLaER1 monocytes (Fig. 5E). Moreover, IL-6 production induced by exogenous M3.1 expression was suppressed in MYXV-infected cells (Fig. 5E), supporting the idea that MYXV may encode additional virulence factors to antagonize the M3.1-mediated NF-κB response.

While we were unable to measure NF-κB induction by M3.1 during MYXV infection, we observed that N4BP1 protein levels were reduced in an M3.1-dependent manner, suggesting that M3.1 targets this pathway during MYXV infection (Fig. 5F, S12B). N4BP1 is cleaved by caspase-8 downstream of death receptors, TLR3, and TLR4 (*29, 31*). Even in the presence of Z-IETD-FMK, a caspase-8 inhibitor, we observed reduced N4BP1 protein levels in cells overexpressing M3.1 or infected with wild-type, but not M3.1-deficient, MYXV (Fig. 5F-G, S12B). Together, our data suggest that M3.1 targets N4BP1 during MYXV infection and that downstream NF-κB activation is suppressed, likely by other virulence factors. Multiple layers of host defenses and poxvirus counterattacks is reminiscent of the “zig-zag” model long proposed to underlie the evolution of ETI in plants (*3*).

## Discussion

Our results support a model in which MYXV employs M3.1 to disable the anti-viral ZAP complex and TBK1-mediated type I IFN signaling (Fig. S10). Notably, we find that disruption of these critical host defense pathways by M3.1 activates a compensatory ETI pathway leading to NF-κB signaling, a response known to restrict poxvirus replication (*18*). We identify N4BP1 as a central player in this ETI pathway. Our data help explain and reconcile previous studies indicating that N4BP1 has two apparently divergent functions: (1) N4BP1 promotes ZAP-mediated virus restriction (*26, 28*), and (2) N4BP1 negatively regulates anti-viral NF-κB responses (*29, 31*). We propose that these two apparently divergent responses can be rationalized by a model in which the anti-viral activity of N4BP1 is “self-guarded” (*17*) by virtue of its ability to inhibit NF-κB signaling. According to this model, targeting of N4BP1 by virulence factors, such as M3.1, triggers a secondary host defense mechanism.

In addition to N4BP1, we show that ZC3H12A and TBK1 are also involved in M3.1-induced NF-κB signaling. Using a luciferase reporter assay, we demonstrate that M3.1 strongly inhibits the NYN ribonuclease activity of ZC3H12A, likely leading to the stabilization of ZC3H12A mRNA targets, such as IL-6. In addition to degrading proinflammatory mRNAs, ZC3H12A has also been reported to degrade viral RNA (*46–48*), suggesting that the ZC3H12A NYN ribonuclease domain may also be self-guarded. In our hands, however, deletion of ZC3H12A alone or in combination with other NYN ribonucleases did not affect MYXV replication (Fig. S13). TBK1, another protein that dually promotes anti-viral responses and negatively regulates NF-κB signaling, is also blocked by M3.1, and its inhibition contributes to M3.1-mediated NF-κB induction. Together with recent work from our group identifying the self-guarded MORC3 pathway (*17*), our results suggest that integration of anti-viral function and immune repression activity into a single protein may be a common feature of mammalian ETI, functioning as an insurance mechanism for critical anti-viral factors. Host proteins originally identified as negative regulators of immune responses may prove to be a rich source for the discovery of ETI pathways, as any negative regulator could potentially function as a pathogen sensor if targeted by a virulence factor.

It is of great interest to investigate whether virulence factors from other pathogens activate the ETI pathway described in this study. Virulence factors from other DNA and RNA viruses have been reported to interfere with ZAP complex activity (*21*). Whether these virulence factors unleash NF-κB signaling remains unknown. Our characterization of M3.1 orthologs indicates that virulence factors from related poxviruses have widely variable effects on NF-κB signaling, suggesting that the poxvirus Bcl-2-like proteins may be rapidly evolving. Comparing the genetic and molecular determinants that drive these diverse activities warrants further study and will likely shed light on the precise mechanisms by which M3.1 and its orthologs interact with their targets.

Importantly, our work establishes an experimental screening approach for the discovery of novel ETI pathways. We anticipate that this approach can be extended to screen additional virulence factors encoded by both viral and bacterial pathogens and will likely provide further insight into the extent and diversity of mammalian ETI.

## Materials and Methods

### Cell culture

BLaER1 cells were cultured in RPMI 1640 medium (Gibco) supplemented with L-glutamine, sodium pyruvate, 100 U/ml penicillin-streptomycin, and 10% (v/v) FBS (Gibco). HEK293T and RK13 cells were cultured in DMEM medium (Gibco) containing the same supplements. BLaER1 cells were transdifferentiated into monocytes for 5-6 days in medium containing 10 ng/ml of human recombinant (hr) IL-3, 10 ng/ml hr-CSF-1 (M-CSF) (both PeproTech), and 100 nM β-Estradiol (Sigma-Aldrich) as previously described (*49*). A total of 1.4 million BLaER1 cells were transdifferentiated per well of a 6-well plate, and 7×10^4^ cells were transdifferentiated per 96 well. BLaER1 cells were a gift from Thomas Graf (CRG, Barcelona, Spain) and Veit Hornung (LMU Munich, Germany). HEK293T and RK13 cells were from the UC Berkeley Cell Culture Facility. All cell lines were routinely tested to be free of mycoplasma contaminations.

### Cell stimulation

Cells were stimulated with the indicated concentrations of LPS-EB Ultrapure from *E. coli* O111:B4 (Invivogen) or hrTNFα (PeproTech). For activation of doxycycline-inducible transgene expression, cells were treated with 1 µg/ml doxycycline hyclate (Sigma-Aldrich) for the indicated time.

### Cloning

A doxycycline-inducible lentiviral vector (pLIX_403, Addgene #41395) was used for transgene expression in BLaER1 cells. Gateway cloning was used to insert transgenes in frame with the vector V5 tag unless otherwise stated. For transient transfection in HEK293T cells, the pGCS vector system (Addgene, Kit #1000000107) was used. The pGCS vector system is based on a pCS2+ backbone that has been modified to be compatible with Gateway cloning and to bear either N-or C-terminal epitope tags. Specifically, pGCS1 (no tag), pGSC-N2 (N-terminal 3xHA tag), pGCS-N3 (N-terminal 3xFLAG tag), and pGCS-C3m (C-terminal 3xFLAG) were used in this study. Gene blocks bearing Gateway attB cloning sites were synthesized by Integrated DNA Technologies and cloned into the Gateway entry vector pDONR201 using Gateway BP Clonase II Enzyme mix (Invitrogen) according to the manufacturer’s instructions. Gateway LR Clonase II Enzyme mix (Invitrogen) was then used to clone genes into the appropriate pGCS vector. Gene mutations were introduced using the Q5-Site Directed Mutagenesis Kit (New England Biolabs) according to the manufacturer’s instructions.

### Lentiviral transduction

Lentivirus was produced in HEK293T cells in 6-well plates. A 90% confluent well was transfected with 1.56 μg of lentiviral vector, 1.17 μg of pd8.9 packaging vector, and 0.468 μg pVSVG using 8 μl Lipofectamine 2000 (Invitrogen) according to the manufacturer’s instructions. After 8-16 hours, the medium was replaced with DMEM medium containing 30% (v/v) FCS and incubated for 24 hours. Viral supernatants were then harvested, centrifugated at 1000 g for 10 minutes, and filtered through a 0.45 μm filter. Following transduction, cells were cultured for 48 hours and then selected with puromycin (Sigma-Aldrich).

### MYXV ORF arrayed screen

A MYXV ORF library in Gateway entry vectors was constructed as described (*12*). Gateway LR Clonase II Enzyme mix (Invitrogen) was used to clone MYXV ORFs, as well as control genes mCherry and adenovirus E4ORF3, into a doxycycline-inducible lentiviral vector (pLIX_403, Addgene #41395). Plasmids were Sanger sequenced by Elim Biopharma to verify sequences.

Lentivirus was produced in HEK293T cells in an arrayed format using 6-well plates as described above. Viral supernatants were harvested and centrifuged at 1000 g for 10 minutes in deep-well 96-well plates. In place of filtering the supernatants, a blasticidin-resistant (Cas9-expressing) BLaER1 monoclone was transduced in medium containing blasticidin (Invivogen) to prevent growth of any remaining HEK293T cells, which were blasticidin-sensitive. After 48 hours, transduced cells were selected with puromycin (Sigma-Aldrich). Successful transductions were achieved for 117 MYXV ORFs (Table S1). BLaER1 B-cells were transdifferentiated into monocytes in a 96-well plate for 5 days as described above and then treated with 1 µg/ml doxycycline hyclate (Sigma-Aldrich) for 24 hours to induce virulence factor expression. Two biological replicates for each MYXV ORF were collected, and prime-seq (*11*) was adapted to prepare libraries for RNA-seq as follows. Cells were washed 1x in PBS and then lysed in TRK lysis buffer (Omega Bio-tek). Lysates were incubated with Proteinase K (0.6 mg/ml) and EDTA (1 mM) for 10 minutes at 50°C. To terminate the reaction, samples were incubated for 5 minutes on ice with PMSF (2 mM). Lysates were combined with 30% PEG beads prepared in house (Thermo Sera-Mag Speed Beads #45152105050250 resuspended in 2 M NaCl, 10 mM Tris-HCl pH 8.0, 1 mM EDTA, 0.01% Igepal CA-630, 0.05% sodium azide, 30% PEG 8000) at a 1:2 ratio of lysate to beads. The beads were washed 2x in 80% ethanol and eluted in DEPC-treated water. Samples were treated with DNase I (Thermo) in the presence of RNasin Plus Ribonuclease Inhibitor (Promega). Reverse transcription into cDNA was performed as described (*11*) in the presence of RNasin Plus Ribonuclease Inhibitor (Promega). During cDNA synthesis, prime-seq uses poly(A) priming, template switching, early barcoding, and UMIs to generate 3′ tagged RNA-seq libraries. Samples from all conditions were then pooled and cleaned up with 30% PEG beads at a ratio of 1:1 lysate to beads. Pooled samples were then treated with Exonuclease I (Thermo). cDNA was then amplified using Terra PCR Direct Polymerase Mix (Takara) for 9 cycles. Libraries were then tagmented (4-5 technical replicates per library) and amplified using Nextera XT DNA Library Preparation Kit (Illumina) according to the manufacturer’s instructions. The following primers were used for library amplification and i7 adaptor addition: 3′ enrichment primer (P5NEXTPT5): AATGATACGGCGACCACCGAGATCTACACTCTTTCCCTACACGACGCTCTTCCGATC T i7 index primer: CAAGCAGAAGACGGCATACGAGAT[i7]GTCTCGTGGGCTCGG Libraries were then pooled and sequenced by the Vincent J. Coates Genomics Sequencing Lab (UC Berkeley) on two Illumina NovaSeq 6000 SP flow cells using 10X sequencing specifications. zUMIs (*50*) was used to demultiplex samples, collapse UMIs, and align reads to human (GRCh38) and myxoma virus (GCF_000843685.1) reference genomes. Differential gene expression analysis was conducted by iterating through each condition and using DESeq2 (1.42.1) to compare gene expression to that of mCherry-expressing control samples. Code for generating figures is available on Github: https://github.com/brennaremick/MYXV_ORF_screen_RNAseq

### RNA-seq

2.8 million BLaER1 monocytes expressing doxycycline-inducible mCherry or M3.1 were treated with 1 µg/ml doxycycline hyclate (Sigma-Aldrich) for 24 hours. Three biological replicates were collected for each genotype and condition. RNA was isolated using the Omega Biotek Total RNA Kit I according to the manufacturer’s instructions. DNA was removed with TURBO DNase (Invitogen) in the presence of RNasin Plus Ribonuclease Inhibitor (Promega), and RNA was isolated with RNAClean XP beads (Beckman Coulter). For Figure 1C-D, library preparation and sequencing were performed by the QB3-Berkeley Genomics core using the KAPA mRNA Hyper Prep kit (Roche KK8581). Libraries were sequenced on an Illumina NovaSeq 6000 S4 flow cell (2×150 bp, 25M paired end reads per sample). For Figure 4B-C, library preparation and sequencing were conducted at Azenta Life Sciences using the NEBNext Ultra II RNA Library Prep Kit for Illumina (New England Biolabs). Libraries were sequenced on an Illumina NovaSeq (2×150 bp, 20M paired end reads per sample). Sequencing quality of fastq files was evaluated with FastQC, and adaptors were trimmed using Cutadapt (1.18). Paired end RNA-seq reads were aligned to the reference genome (GRCh38) using STAR (2.7.1a). Fragment counts were quantified using featureCounts from the Subread (2.0.3) package, and differential gene expression analysis was conducted using DESeq2 (1.42.1). Gene set enrichment analysis was conducted using clusterProfiler (4.10.1). The Molecular Signatures Database (MSigDB) hallmark gene sets were downloaded using the R package msigdbr (7.5.1). Code for generating figures is available on Github:

https://github.com/brennaremick/MYXV_M003-1_RNAseq_1

https://github.com/brennaremick/MYXV_M003-1_RNAseq_2

### Cytokine quantification

Cytokine secretion was quantified by ELISA of cell-free supernatants following the manufacturer’s instructions (human IL-6: BD 555220).

### Immunoprecipitation mass spectrometry

Ten 15-cm plates per condition were seeded with 4 million HEK293T cells. After 24 hours, cells were transfected with 5 μg M3.1-3xFLAG per plate. 48 hours post transfection, cells were treated with 10 ng/ml hrTNFα (PeproTech) or media control for 8 hours and then harvested by scraping in PBS. Cell pellets were flash frozen in liquid nitrogen. Cells were lysed in lysis buffer (40 mM HEPES pH 7.5, 150 mM NaCl, 0.2% NP-40, cOmplete EDTA-free protease inhibitor cocktail tablets (Sigma Aldrich)) for 60 minutes at 4°C. Lysates were spun at 21,000 g for 30 minutes, and the supernatant was added to 90 μl of prewashed ANTI-FLAG® M2 Affinity Agarose Gel slurry (Sigma-Aldrich, A2220). After 1.5 hours of rocking, the beads were spun down and washed 5x with lysis buffer. Beads were then washed 2x in PBS with 0.2% NP-40.

Proteins were eluted with 500 μg/ml 3xFLAG peptide (Millipore Cat#F4799) in PBS with 0.2% NP-40. Elutions were precipitated in 20% final concentration of trichloroacetic acid on ice overnight. Precipitations were spun at 21,000 g for 10 minutes and washed 3x in ice cold acetone and dried. The pellets were solubilized in 8 M urea 100 mM Tris pH 8.5, treated with TCEP and iodoacetamide, and digested overnight with trypsin (Promega). Samples were analyzed by Multidimensional Protein Identification Technology (MudPIT) at the Vincent J. Coates Proteomics/Mass Spectrometry Laboratory (UC Berkeley). M3.1-interacting proteins were identified by CompPASS analysis (*51*) by comparing the samples to over 70 similarly performed anti-FLAG immunoprecipitations from HEK293T cells. Total spectral counts were normalized to 1000 bait counts, and results with a Z-score greater than 5 were plotted. See Data S1 for CompPASS output.

### Coimmunoprecipitation

Cells were lysed in lysis buffer (40 mM HEPES pH 7.5, 150 mM NaCl, 1% NP-40, Pierce Protease and Phosphatase Inhibitor EDTA-free mini tablets (Thermo)) for 30 minutes on ice. Lysates were clarified by centrifugation for 30 minutes at 4°C, and 5% of clarified lysate was removed as an input. Remaining sample was added to 20 μL of washed ANTI-FLAG® M2 Affinity Agarose Gel slurry (Sigma-Aldrich, A2220) and rotated for 1-2 hours at 4°C. Beads were washed four times and eluted with Laemmli buffer by boiling for 7 minutes. Samples were then analyzed by immunoblot.

### Immunoblotting and antibodies

Whole cell lysates were prepared by lysing cells in RIPA buffer (150 mM NaCl, 5 mM EDTA pH 8.0, 50 mM Tris-HCl pH 8.0, 1% NP-40, 0.5% sodium deoxycholate, 0.1% SDS) supplemented with Pierce Protease and Phosphatase Inhibitor EDTA-free mini tablets (Thermo) for 20 minutes on ice. Lysates were clarified by centrifugation for 20 minutes at 4°C. Laemmli buffer was added to a final concentration of 1x and lysates were boiled at 95°C for 7 minutes. Samples were run on 4-12% Bis-Tris protein gels (Invitrogen) and then transferred to Immobilon-FL PVDF membranes (Millipore Sigma). Membranes were blocked with Li-Cor Odyssey blocking buffer and then incubated with primary antibodies overnight at 4°C. Following incubation with appropriate secondary antibodies, blots were imaged using the Li-Cor Odyssey platform.

Antibodies used were:

Phospho-NF-κB p65 (Ser536) (93H1) (Cell Signaling, #3033)

NF-κB p65 (L8F6) (Cell Signaling, #6956)

IκBα (Cell Signaling, #9242)

V5-Tag (D3H8Q) (Cell Signaling, #13202)

Actin (Santa Cruz, sc-47778)

TRIM25 (Abcam, ab167154)

ZC3HAV1/ZAP (Proteintech, 16820-1-AP)

N4BP1 (Thermo Fisher/Bethyl Laboratories, #A304-628A-T)

KHNYN (Santa Cruz, sc-514168)

ZC3H12A (GeneTex, GTX110807)

TBK1/NAK (D1B4) (Cell Signaling, #3504)

Phospho-TBK1/NAK (Ser172) (D52C2) (Cell Signaling, #5483)

FLAG M2 (Sigma-Aldrich, F3165)

FLAG (D6W5B) (Cell Signaling, #14793)

TANK (Cell Signaling, #2141)

IKKε (Cell Signaling, #2690)

HA-tag (C29F4) (Cell Signaling, #3724)

IRDye 800 Donkey anti-Mouse IgG Secondary Antibody (Li-Cor, 926-32212) Alexa Fluor 680 Goat anti-Rabbit IgG Secondary Antibody (Invitrogen, A-21109)

### Gene disruption by Cas9-RNP nucleofection

Polyclonal gene-deficient BLaER1 cells and HEK293T cells were generated as follows. Two gRNAs per gene were designed to target early coding exon(s) of the respective gene using ChopChop (*52*). BLaER1 cells expressing doxycycline-inducible mCherry or M3.1 or HEK293T cells were harvested and washed 1x in PBS. 2 million cells were nucleofected with Alt-R® S.p. Cas9 Nuclease V3 (IDT) complexed with two gRNAs per gene (Synthego, sgRNA EZ Kit) and Alt-R® Cas9 Electroporation Enhancer (IDT) in Lonza P3 buffer (Lonza, V4XP-3032) as described (*34*). Nucleofection was performed with a Lonza 4DNucleofector Core Unit (AAF-1002B) using the program CM-137. Negative Control sgRNA (mod) #1 (Synthego) was used as a non-targeting control (NTC). Knockout efficiency was evaluated by immunoblot (Fig. S5-6).

Deletion of multiple genes was performed sequentially. gRNA sequences used (PAM is highlighted in bold):

*N4BP1* gRNA 1: GACCTTGCATCAGTAACCGA**AGG**

*N4BP1* gRNA 2: ACAGGCCCTCGATACGGCCG**CGG**

*KHNYN* gRNA 1: ATGGAAACGAGGCGCCCGAG**GGG**

*KHNYN* gRNA 2: GAGCTCCCCCTAGTGACGGC**AGG**

*ZC3H12A* gRNA 1: GAATCGGCACTTGATCCCAT**AGG**

*ZC3H12A* gRNA 2: GGTCATCGATGGGAGCAACG**TGG**

*TRIM25* gRNA 1: GTCGTGCCTGAATGAGACGT**GGG**

*TRIM25* gRNA 2: GCGGCGCAACAGGTCGCGAA**CGG**

*ZC3HAV1* (ZAP) gRNA 1: CGGACTGCGAATAGTTGCAC**CGG**

*ZC3HAV1* (ZAP) gRNA 2: AAAATCCTGTGCGCCCACGG**GGG**

*TBK1* gRNA 1: GGTAGTCCATAGGCATTAGA**AGG**

*TBK1* gRNA 2: AAATATCATGCGTGTTATAG**GGG**

*IKBKE* (IKKε) gRNA 1: TGCATCGCGACATCAAGCCG**GGG**

*IKBKE* (IKKε) gRNA 2: GCCCCAGCAAAAAGCGTTCG**GGG**

*TANK* gRNA 1: GCAGAGAATACGTGAACAAC**AGG**

*TANK* gRNA 2: CCACAAGATAAAGTGATTTC**AGG**

### MYXV preparation

Recombinant viruses were generated by homologous recombination. The DNA fragment containing a reporter gene that replaced the target ORF and appropriate flanking sequences for the homologous recombination was synthesized and cloned in the pUC57 plasmid by GenScript. To generate the vMyx-ΔM003.1 virus, the M3.1 ORF was replaced with a GFP expression cassette (driven by a poxvirus synthetic early/late promoter) flanked by the M-T2 (partial) and M003.2 sequences. RK13 cells were infected with the wild-type MYXV-Lau strain for one hour, and then the unbound virus was removed. Cells were then transfected with the recombination plasmid using Effectene transfection reagent (Qiagen). GFP expression from the recombination plasmid was monitored using a Leica fluorescence microscope. The cells were scraped and collected with media 48 hours post-infection and stored at –80°C until processed. The samples were freeze-thawed at –80°C and 37°C three times and sonicated with a Cup Horn Sonicator for 1 minute to release the viruses from the infected cells. The samples were serially diluted with media, plated on RK13 cells, and infected for 1 hour. Media was removed and layered with 1% low-melting agarose diluted with the media. After 48 hours of infection, fluorescent foci were selected using the microscope, and foci were picked using pipette tips and diluted in media.

Subsequently, multiple rounds of foci purification were performed until the pure foci of only recombinant viruses were isolated. The virus was amplified from a single foci in RK13 cells, and the purity of the virus was confirmed by PCR using appropriate primers. The construction of wild-type MYXV that expresses GFP under the control of a poxvirus synthetic early/late promoter (vMyx-GFP) was described previously (*53*). All the viruses were amplified after infecting RK13 cells grown in multiple T150 cell culture dishes. Viruses were then purified by centrifugation through a sucrose cushion as described previously (*54*).

### Myxoma virus infections

Transdifferentiated BLaER1 monocytes and HEK293T cells were infected with MYXV at the indicated MOI. For virus replication assays, cells were infected for 1 hour, and then unbound virus was removed and replaced with fresh media. Cells were scraped and collected with media at various times post-infection and stored at −80°C until processed. The samples were freeze-thawed at −80°C and 37°C three times to release the virus from infected cells. The virus was titrated onto confluent RK13 monolayers by serial dilution in triplicate. After 48 hours of infection, fluorescent foci were quantified using a Zeiss Cell Discoverer 7 at the Cancer Research Laboratory (CRL) Molecular Imaging Center (UC Berkeley) and used to calculate the virus titer.

### AlphaFold 3 structure predictions

Amino acid sequences for proteins of interest were obtained from NCBI. The AlphaFold 3 (*39*) server (https://alphafoldserver.com/) was used to generate predicted structures of the following complexes:

MYXV M3.1 (QCO69335.1), N4BP1 (NP_694574.3)

MYXV M3.1 (QCO69335.1), ZC3H12A (NP_001310479.1)

MYXV M3.1 (QCO69335.1), TRIM25 (NP_005073.2)

Output models were visualized in ChimeraX (1.8). Predicted Aligned Error (PAE) plots were visualized in PAE Viewer (*55*).

### NF-κB and IFNΔ luciferase assays

HEK293T cells were plated in white-walled 96-well plates and incubated overnight. Using Lipofectamine 2000 (Invitrogen), HEK293T cells were co-transfected with the indicated constructs, reporter constructs (50 ng) expressing firefly luciferase under control of the *ELAM* promoter (NF-κB reporter) or the *IFNB1* promoter, and a *Renilla* luciferase-expressing construct (10 ng) to control for transfection efficiency. After 24-36 hours, luciferase activity was measured using the Dual-Glo Luciferase Assay System (Promega) according to the manufacturer’s instructions. To account for transfection efficiency, the firefly luciferase values were divided by *Renilla* luciferase values for each well.

### ZC3H12A ribonuclease activity luciferase assay

The pmirGLO Dual-Luciferase expression vector (Promega) was used to measure ZC3H12A ribonuclease activity. A ZC3H12A target, the human *IL-6* 3’UTR, was cloned downstream of the firefly luciferase gene. Using Lipofectamine 2000 (Invitrogen), HEK293T cells were co-transfected with the pmirGLO-IL6 3’UTR construct (50 ng) and constructs encoding ZC3H12A, M3.1, and/or mCherry controls. After 24 hours, luciferase activity was measured using the Dual-Glo Luciferase Assay System (Promega) according to the manufacturer’s instructions. The internal *Renilla* luciferase control was used to normalize for transfection efficiency.

### Phylogenetic analysis

Position-specific iterated (PSI)-BLAST was used to identify M3.1 orthologs. Orthologs that were able to be expressed in HEK293T cells were selected for further analysis. VACV A52, a Bcl-2-like protein reported to activate MAPK signaling, was chosen as an outgroup (*19, 42*). The following protein sequences were downloaded from NCBI:

Myxoma virus M3.1 (QCO69335.1)

Rabbit fibroma virus (NP_051888.1)

Lumpy skin disease virus (AYV61288.1)

Swinepox virus (QQG31492.1)

Cowpox virus (NP_619989.1)

Vaccinia virus (Western Reserve strain) B14 (UZL86927.1)

Vaccinia virus (Western Reserve strain) A52 (YP_233060.1)

Above sequences were aligned on Phylogeny.fr with MUSCLE using the default settings. Maximum likelihood phylogenetic trees were generated with PhyML using 100 bootstrap replicates.

### Statistical analysis

Statistical analysis was performed in GraphPad Prism 10. Statistical tests are indicated in the figure legends. Each data point is from an independent experiment and represents the average of 2-3 technical replicates.

### Data availability

Raw and processed RNA-seq data is deposited at NCBI Gene Expression Omnibus: GSE288433, GSE287860, GSE288000

### Source code

Code for RNA-seq analysis and figure generation is available on Github:

https://github.com/brennaremick/MYXV_ORF_screen_RNAseq

https://github.com/brennaremick/MYXV_M003-1_RNAseq_1

https://github.com/brennaremick/MYXV_M003-1_RNAseq_2

## Acknowledgments

We thank members of the Vance, Gaidt, Barton, Coscoy, and Ohainle labs for advice and discussions; M. Ohainle, G. Liu, and T. Zhang for comments on the manuscript; S. McWhirter for the TBK1 plasmids; R. Chavez for technical support; A. Janjic for advice with the prime-seq protocol; and K. Heydari, M. Delcroix, and H. Dhaliwal in the CRL Flow Cytometry Facility, L. Kohlstaedt in the Vincent J. Coates Proteomics/Mass Spectrometry Laboratory, H. Aaron in the CRL Molecular Imaging Center, and the QB3 Genomics facilities (all UC Berkeley) for technical advice and assistance.

## Author contributions

Conceptualization: BCR, MMG, REV; Investigation: BCR, JQM, AGM, ADG, MMR, MMG; Data analysis: BCR, JQM, AGM, AW; Resources: MMR, GM, MR, MMG; Funding acquisition: REV; Supervision: MMG, REV; Writing: BCR, MMG, REV.

## Supplemental Figures

**Fig. S1.**
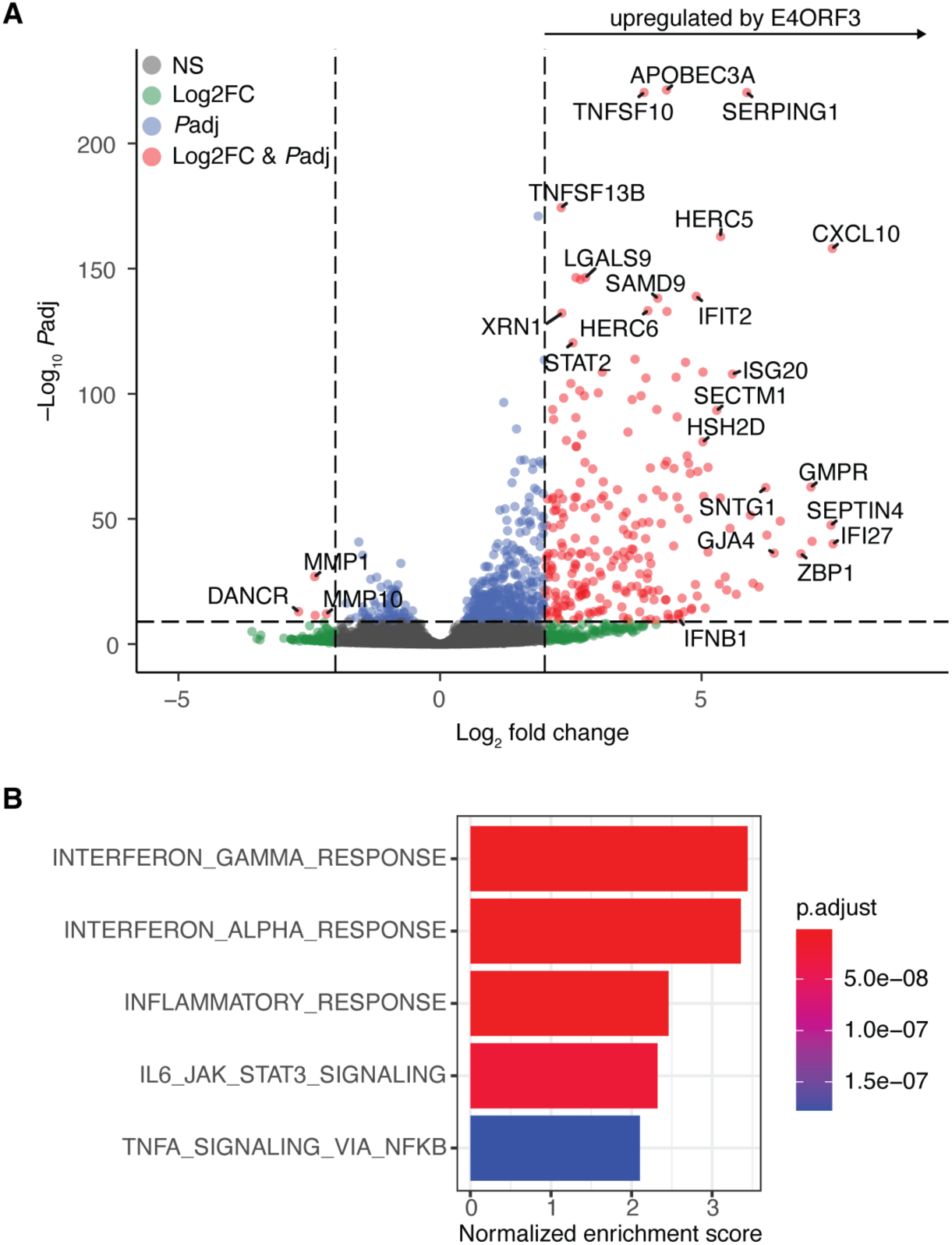
Detection of adenovirus E4ORF3-mediated type I IFN induction validates arrayed virulence factor screening approach. (**A**) Volcano plot from the screen of differentially expressed genes in BLaER1 monocytes expressing E4ORF3 for 24 hours compared to mCherry. Dashed lines indicate a log_2_FC cutoff of 2 and a *P*adj cutoff of 10e-10. **(B)** MSigDB hallmark gene sets enriched in E4ORF3-expressing BLaER1 monocytes in (A).

**Fig. S2.**
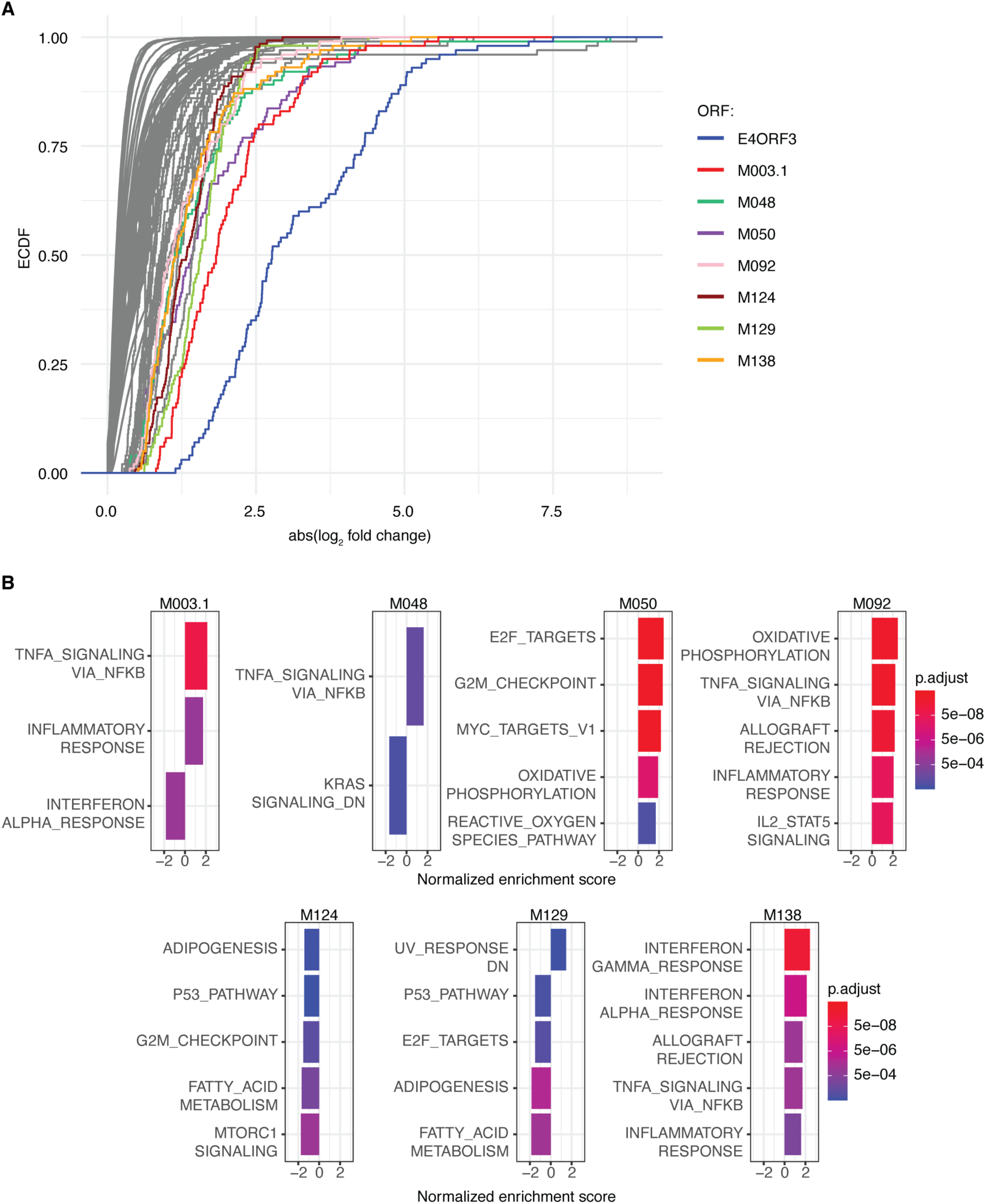
Arrayed screening of the MYXV ORFeome identifies virulence factors that induce host transcriptional responses. (**A**) Differentially expressed genes were identified for each ectopically expressed ORF using DESeq2, with comparisons made against mCherry-expressing control samples. The empirical cumulative distribution function (ECDF) of the absolute fold change was plotted for the top 100 genes with the lowest *P*adj values for each ORF. Colored lines represent specific ORFs that exhibit a distinct shift in the ECDF curve, suggesting a notable transcriptional response induced by these ORFs. (B) MSigDB hallmark gene sets enriched in BLaER1 monocytes expressing the indicated MYXV ORF.

**Fig. S3.**
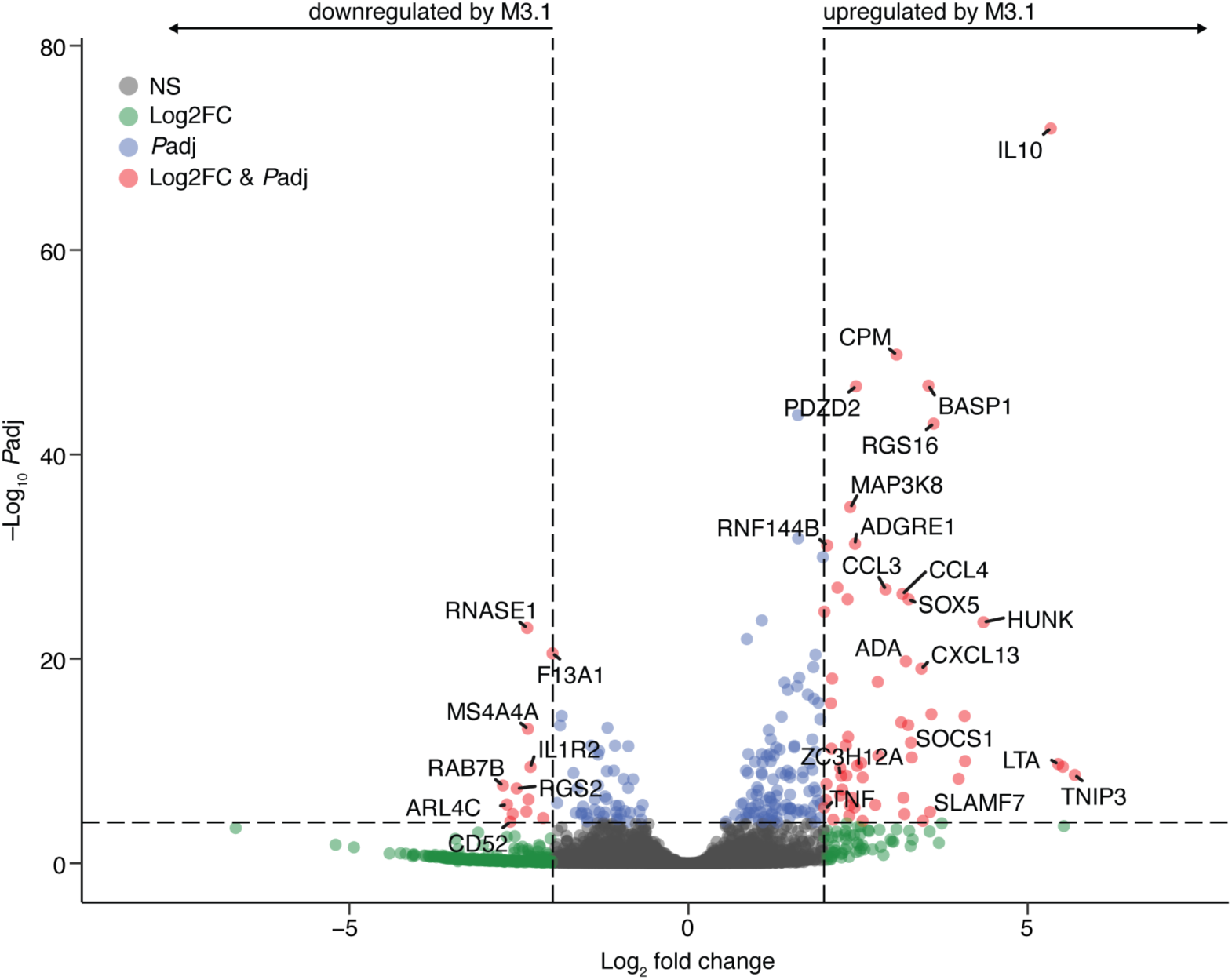
MYXV M3.1 induces a transcriptional response in BLaER1 monocytes. Volcano plot from the screen of differentially expressed genes in BLaER1 monocytes expressing M3.1 (2 biological replicates) for 24 hours compared to mCherry. Dashed lines indicate a log_2_FC cutoff of 2 and a *P*adj cutoff of 10e-5.

**Fig. S4.**
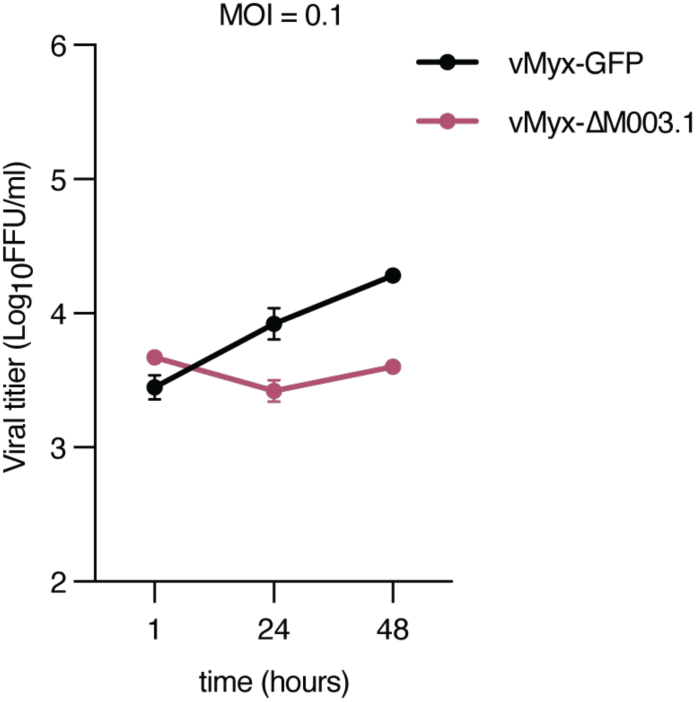
MYXV replicates poorly in BLaER1 monocytes. BLaER1 monocytes were infected with MYXV with an MOI of 0.1. Viral progeny were quantified at the indicated time points.

**Fig. S5.**
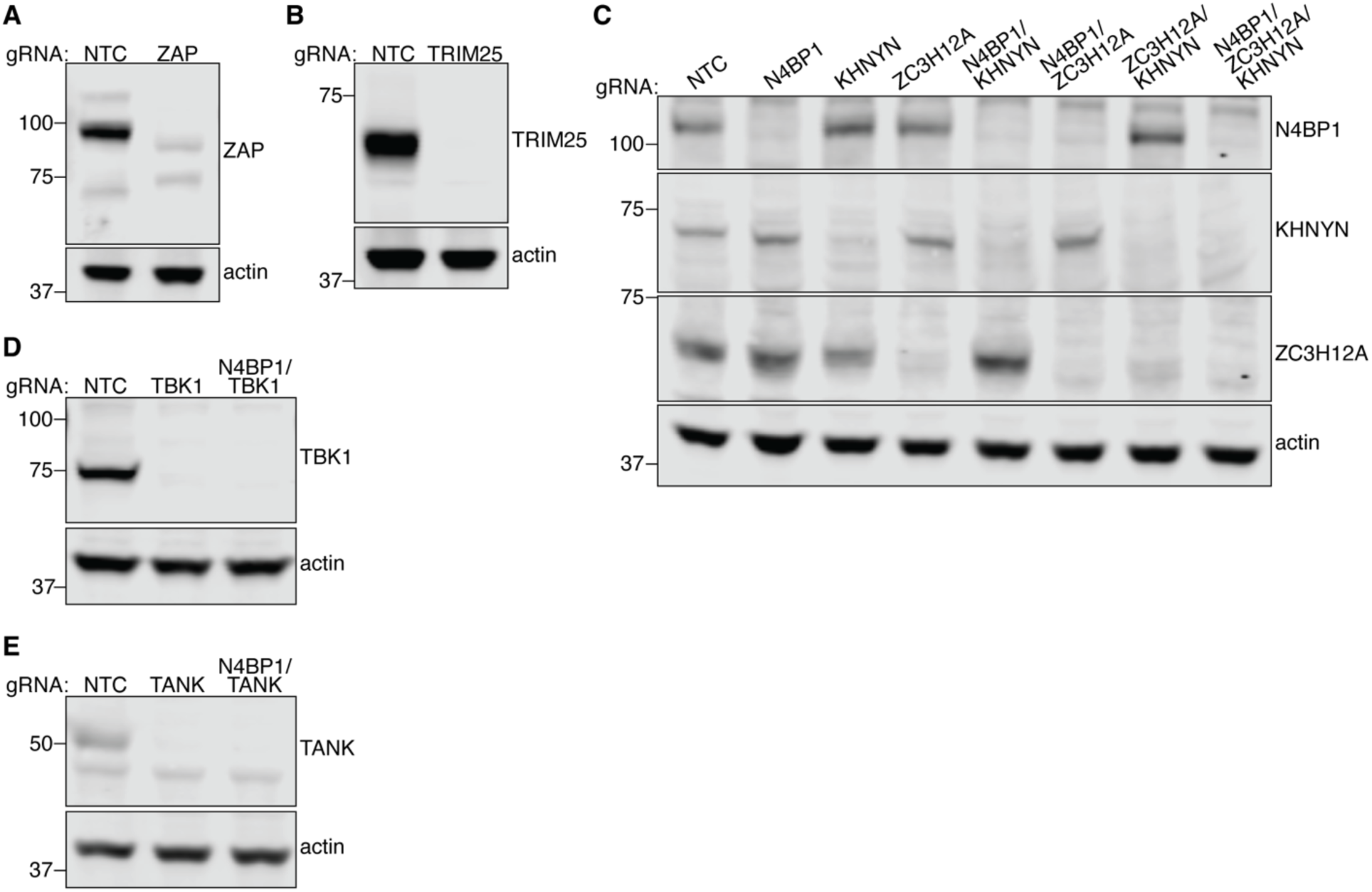
Gene knockout by Cas9-RNP nucleofection of HEK293T cells. HEK293T cells were nucleofected with Cas9 and two guides per gene. Knockout efficiency in the polyclonal population was assessed by immunoblotting for ZAP **(A)**, TRIM25 **(B)**, NYN ribonucleases N4BP1, KHNYN, and ZC3H12A **(C)**, TBK1 **(D)**, and TANK **(E)**. Double or triple knockouts were generated sequentially.

**Fig. S6.**
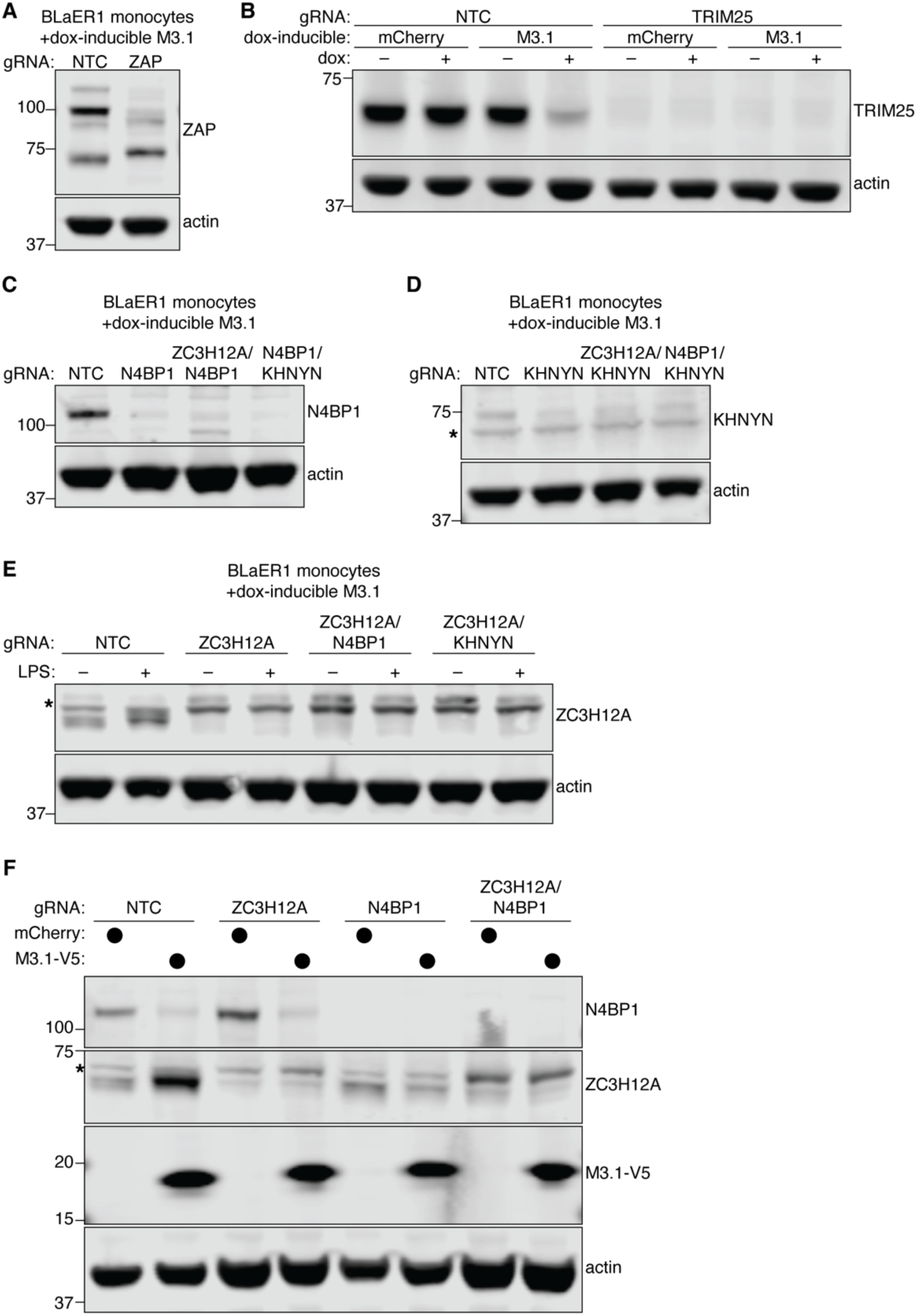
Gene knockout by Cas9-RNP nucleofection of BLaER1 cells. BLaER1 B-cells expressing doxycycline-inducible mCherry or M3.1 were nucleofected with Cas9 and two guides per gene. B-cells were transdifferentiated for 5 days into monocytes, and then knockout efficiency in the polyclonal population was assessed by immunoblotting for ZAP **(A)**, TRIM25 **(B)**, N4BP1 **(C)**, KHNYN **(D)**, and ZC3H12A **(E)**. Since ZC3H12A is an NF-κB-inducible gene, monocytes were stimulated with 200 ng/ml LPS overnight to boost ZC3H12A expression in **(E)**. **(F)** Knockout efficiency and M3.1 expression were verified for RNA-seq samples treated with doxycycline for 24 hours. Double knockouts were generated sequentially. * Indicates non-specific band.

**Fig. S7.**
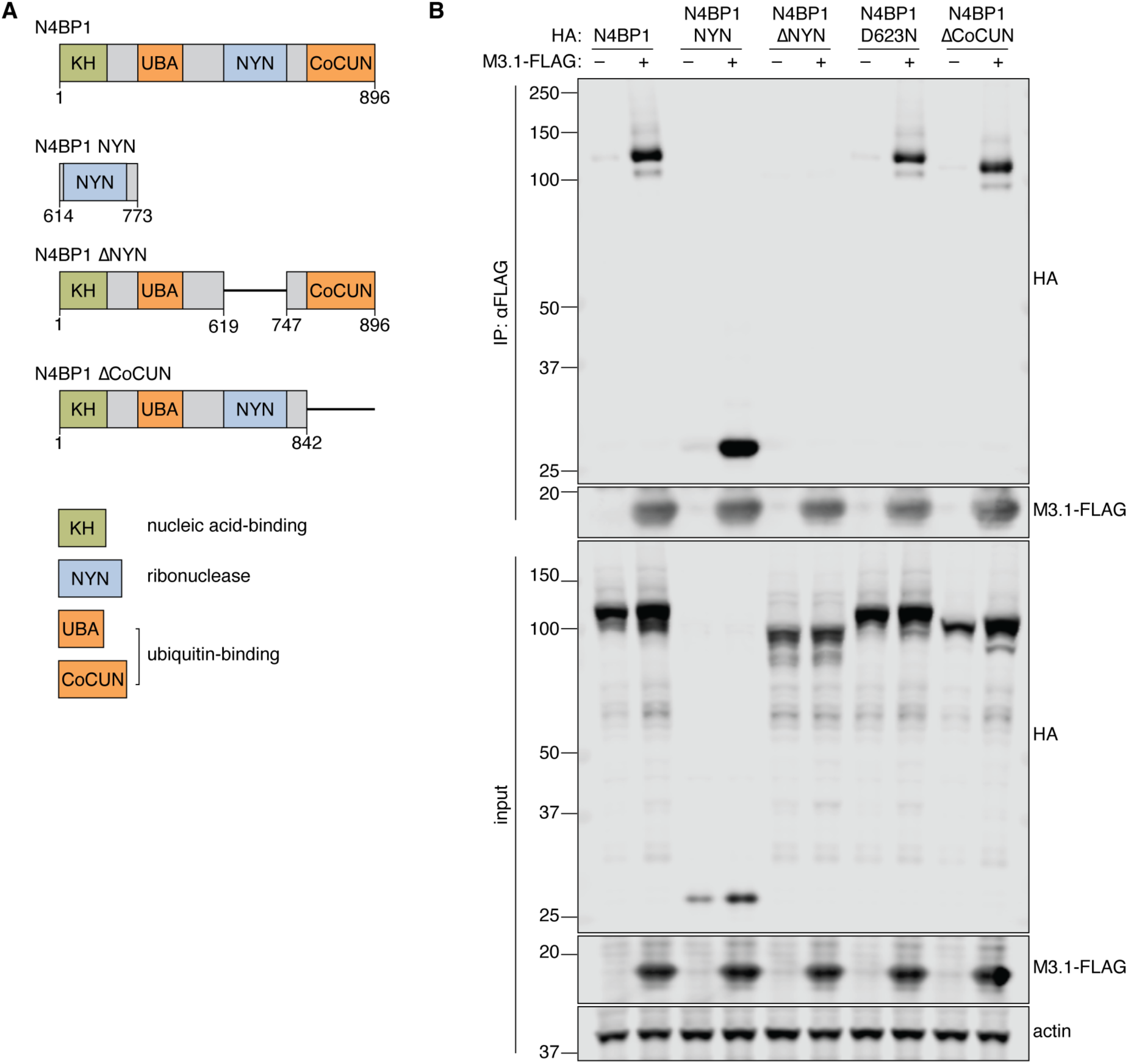
The NYN ribonuclease domain of N4BP1 is necessary and sufficient to interact with M3.1. **(A)** Schematic of N4BP1 domain mutant constructs. KH: K Homology; UBA: Ubiquitin-associated; NYN: N4BP1, YacP-like Nuclease; CoCUN: Cousin of CUBAN. **(B)** Constructs encoding HA-tagged full length N4BP1, N4BP1 NYN domain only, N4BP1ΔNYN, N4BP1 ribonuclease-dead (D623N), and N4BP1ΔCoCUN were expressed in HEK293T cells with M3.1-FLAG or control. M3.1-FLAG was immunoprecipitated and the interaction with N4BP1 variants was assessed by immunoblotting.

**Fig. S8.**
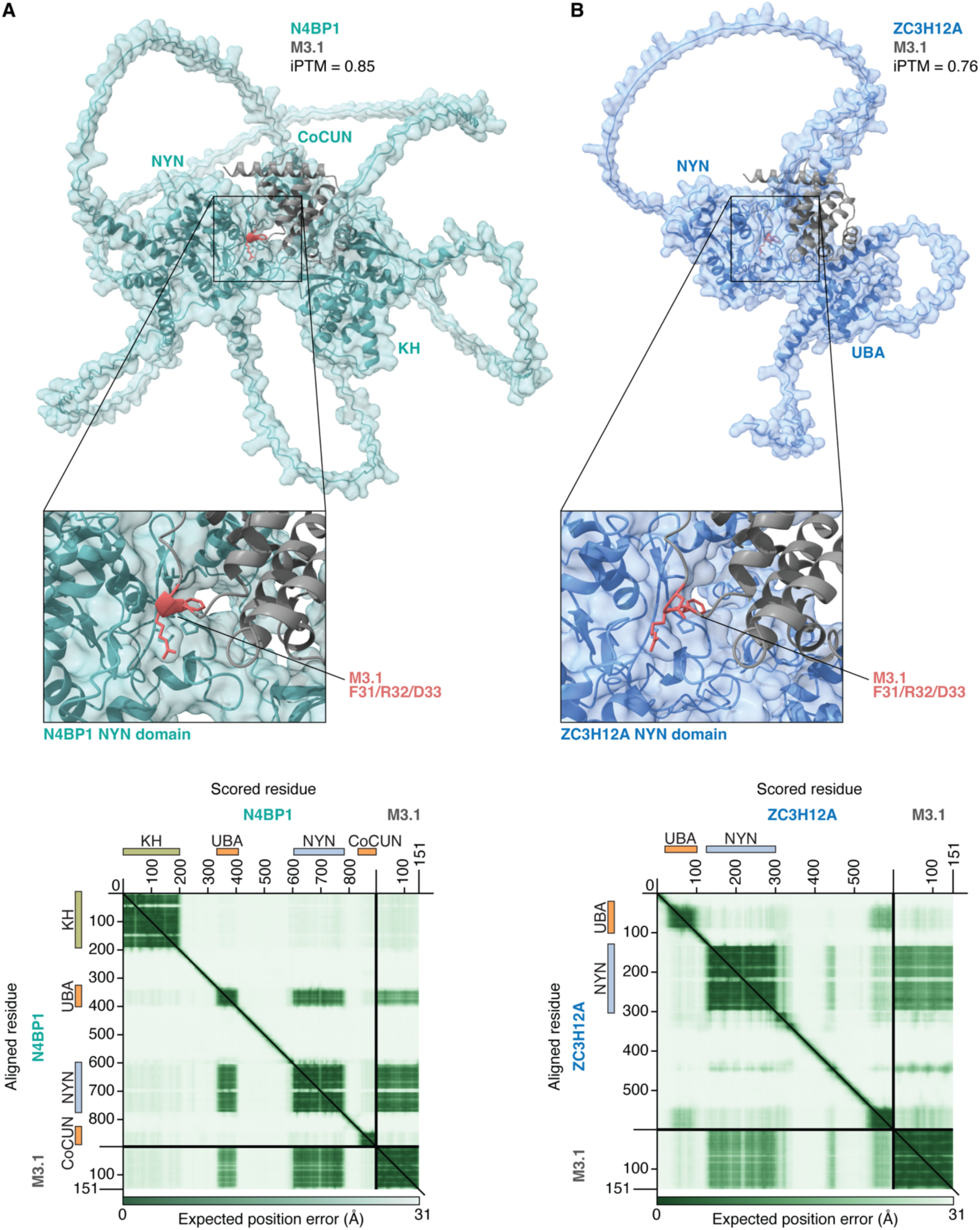
AlphaFold 3-predicted structures of M3.1 with N4BP1 and ZC3H12A. AlphaFold 3 was used to generate predicted structures of the complex of M3.1 with N4BP1 **(A)** and with ZC3H12A **(B)**. Corresponding PAE plots are shown.

**Fig. S9.**
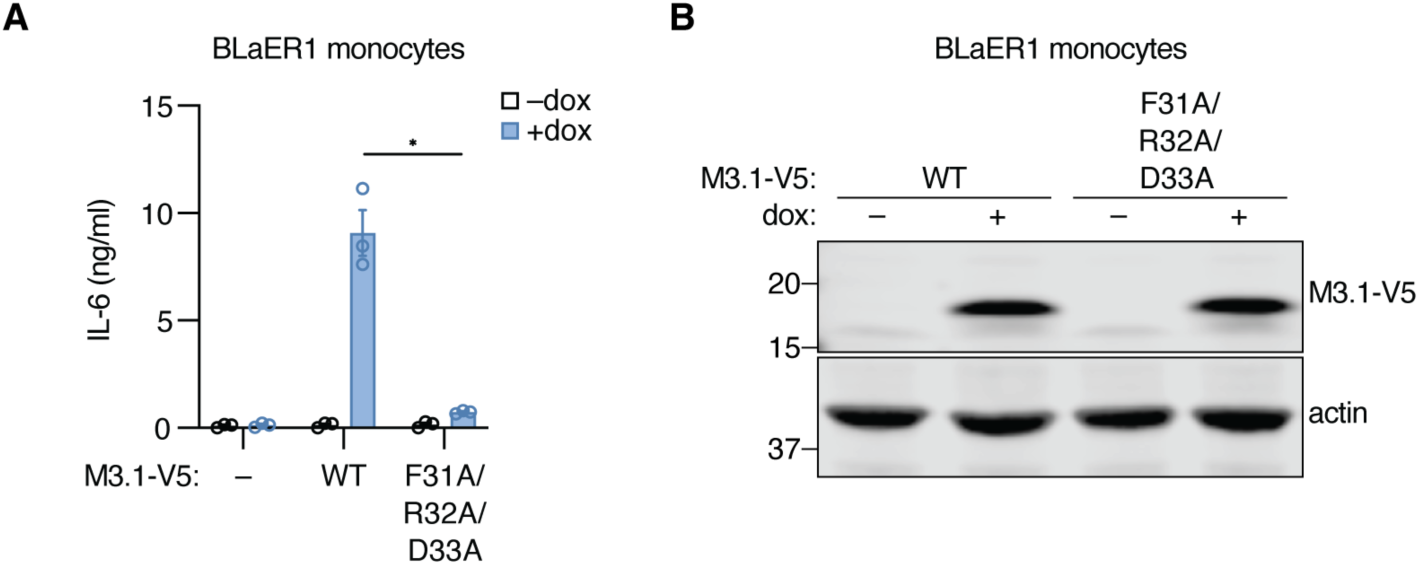
The interaction between M3.1 and NYN ribonucleases is required for M3.1-mediated NF-κB signaling in BLaER1 monocytes. **(A)** Secreted IL-6 from BLaER1 monocytes treated with doxycycline for 24 hours to induce expression of mCherry (–), M3.1 (WT), or M3.1 F31A/R32A/D33A. Data are mean ± SEM of three independent experiments. **(B)** Lysates from (A) were immunoblotted as indicated. * *P* < 0.05; tested by unpaired t-test with Welch’s correction.

**Fig. S10.**
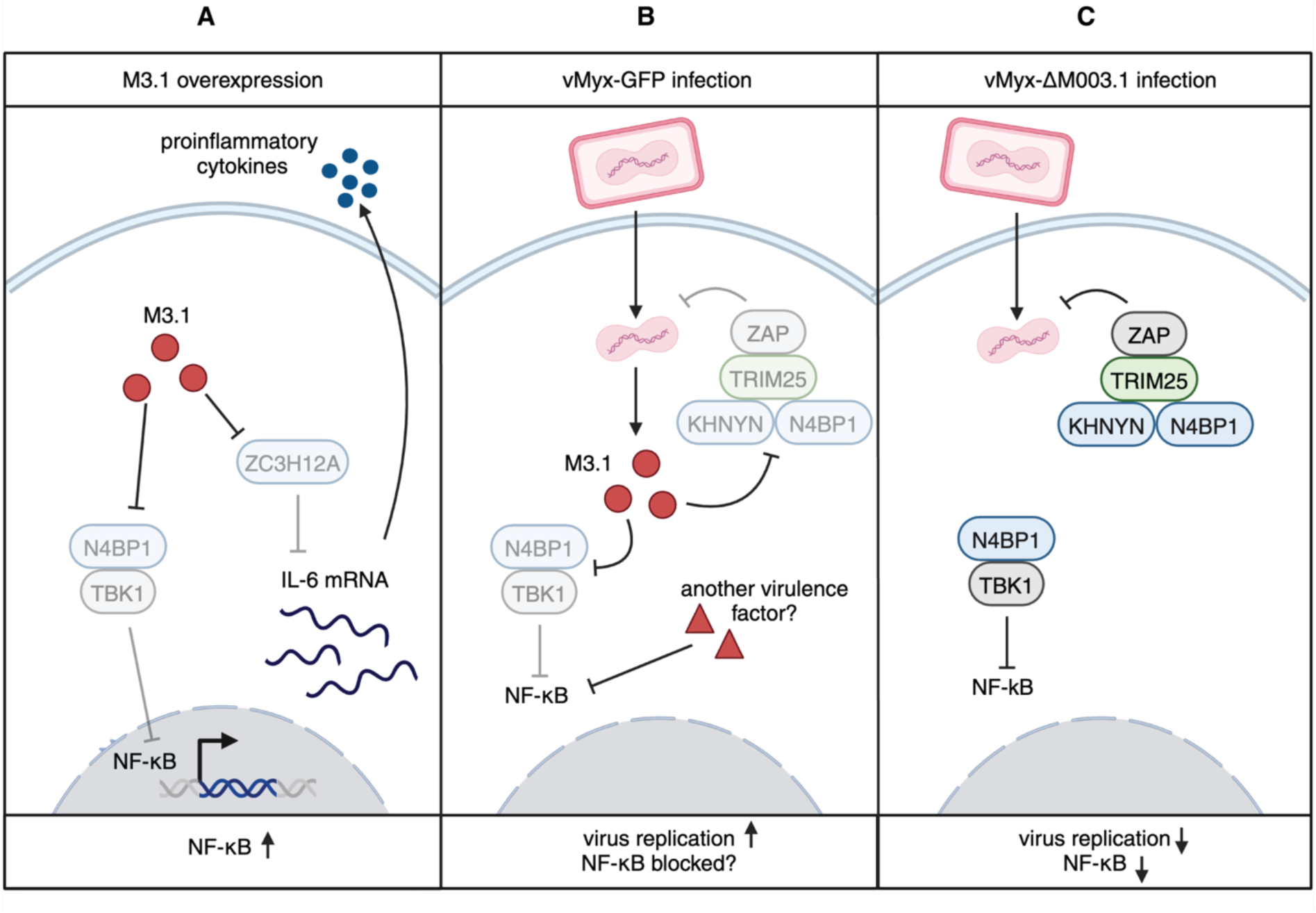
Model for ETI sensing of MYXV M3.1. (**A**) Ectopic expression of M3.1 induces proinflammatory cytokine production by blocking negative regulators of NF-κB signaling N4BP1, ZC3H12A, and TBK1. (B) During MYXV infection, M3.1 blocks the anti-viral ZAP complex, enabling viral replication. M3.1 targets N4BP1 during MYXV infection, but we do not observe NF-κB signaling, possible due to the activity of other virulence factors that suppress NF-κB. (C) M3.1-deficient MYXV is restricted by the anti-viral ZAP complex consisting of ZAP, the E3 ubiquitin ligase TRIM25, and NYN ribonucleases N4BP1 and KHNYN. Created with BioRender.com.

**Fig. S11.**
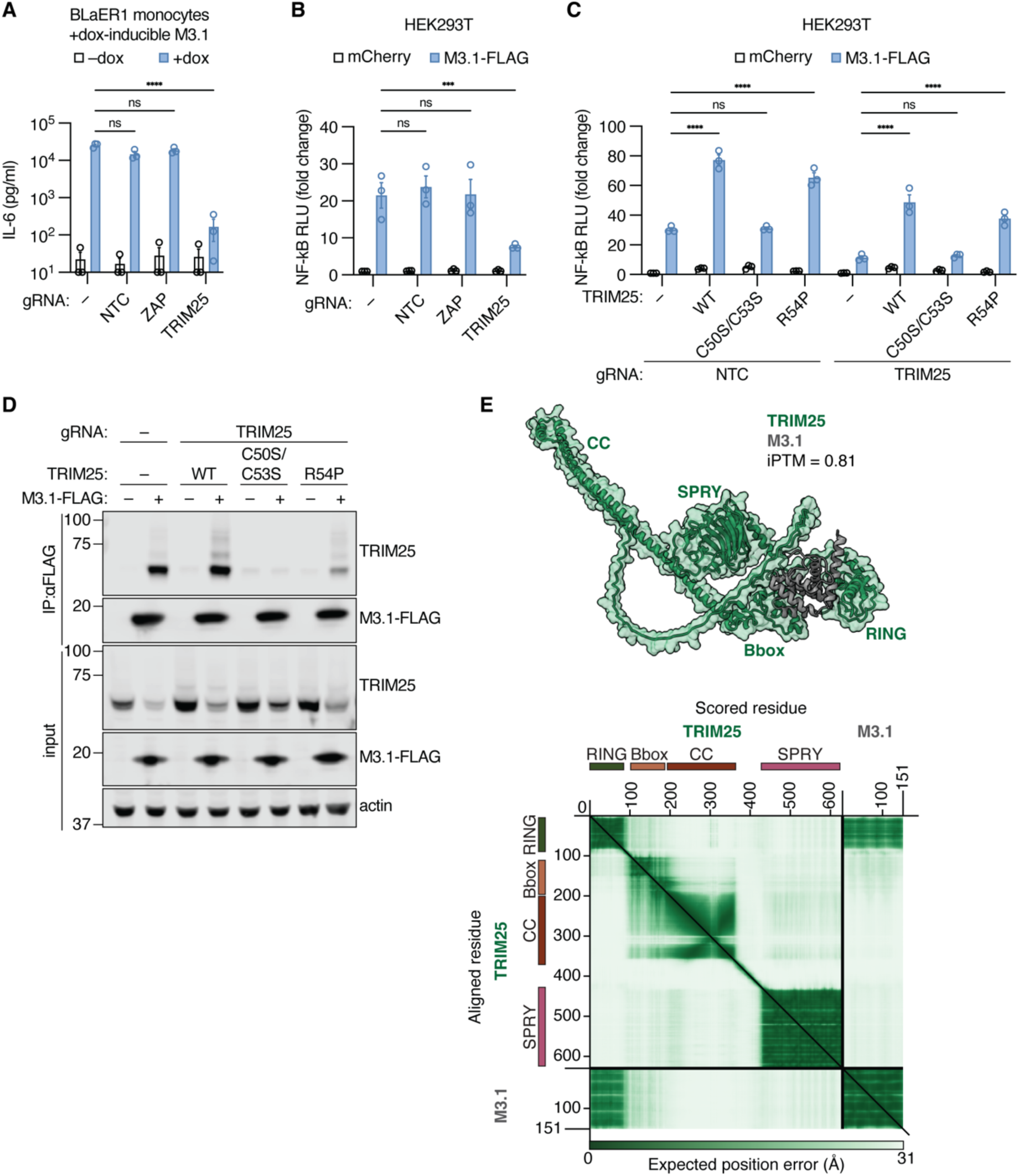
TRIM25 promotes M3.1-mediated NF-κB signaling. (**A**) BLaER1 monocytes expressing doxycycline-inducible M3.1 were nucleofected with Cas9 and two gRNAs targeting the indicated gene. Cells were treated with doxycycline for 24 hours to induce expression of M3.1, and IL-6 production was measured by ELISA. Data are mean ± SEM of three independent experiments. **(B)** Cas9-RNP nucleofection was used to disrupt the indicated genes in HEK293T cells, and NF-κB induction by M3.1 was measured using a luciferase reporter. Data are mean ± SEM of three independent experiments. **(C)** A luciferase reporter was used to measure NF-κB induction in TRIM25-deficient or control HEK293T cells co-expressing M3.1 and TRIM25 variants. Data are mean ± SEM of three independent experiments. **(D)** M3.1-FLAG was immunoprecipitated from WT and TRIM25-deficient HEK293T cells expressing TRIM25 variants. The interaction between M3.1 and TRIM25 was assessed by immunoblotting. **(E)** AlphaFold 3 was used to generate the predicted structure of the complex of M3.1 and TRIM25. Corresponding PAE plot is shown. RING: Really Interesting New Gene; CC: Coiled-Coil; SPRY: SPla and the RYanodine Receptor. *** *P* < 0.001; **** *P* < 0.0001; ns = not significant, tested by 2-way ANOVA with Šídák’s post-hoc test (A-C). Statistical testing for (A) was performed on log-normalized data.

**Fig. S12.**
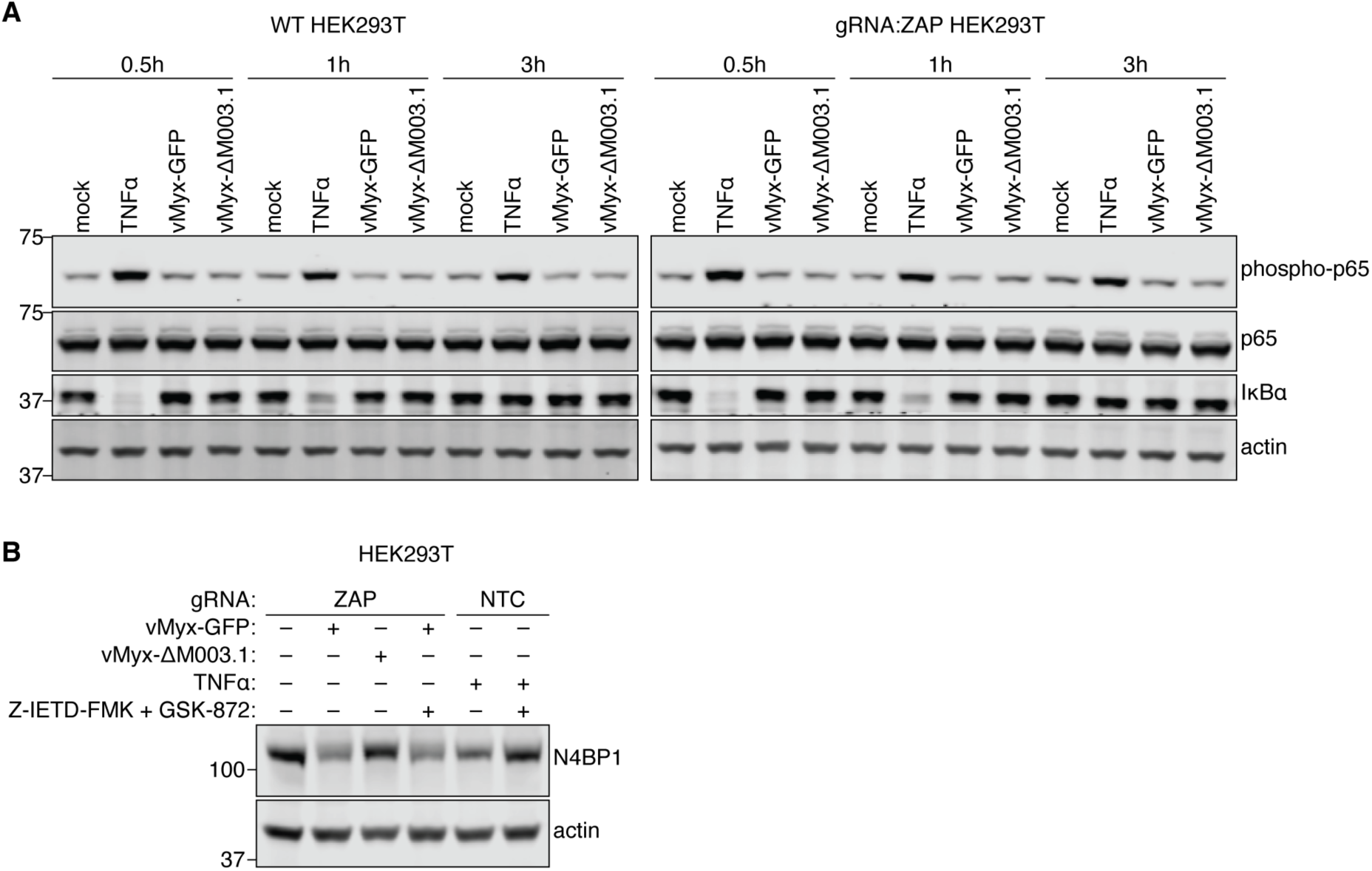
MYXV does not activate NF-κB signaling in HEK293T cells. (**A**) HEK293T cells were either stimulated with TNFα (50 ng/ml) or infected with MYXV (MOI = 3) for the indicated times. Lysates were immunoblotted as indicated. **(B)** HEK293T cells were either stimulated with TNFα (50 ng/ml) or infected with MYXV (MOI = 1) for 24 hours in the presence of caspase-8 (10 μM Z-IETD-FMK) and RIPK3 (3 μM GSK-872) inhibition or DMSO control. Lysates were immunoblotted as indicated.

**Fig. S13.**
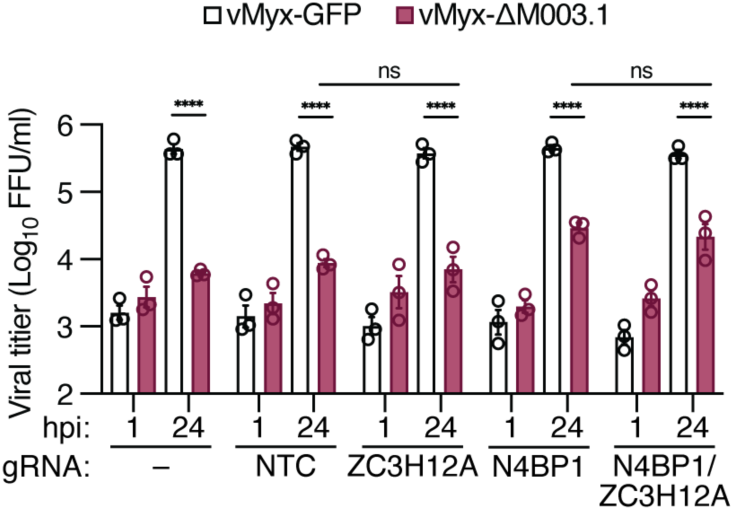
ZC3H12A does not restrict MYXV. MYXV replication in HEK293T cells infected with an MOI of 1. Viral progeny were quantified at the indicated time points. Data are mean ± SEM of 3 independent experiments. **** *P* < 0.0001; ns = not significant, tested by 3-way ANOVA with Tukey’s post-hoc test.

